# A comprehensive characterization of the phospholipid and cholesterol composition of the uncinate fasciculus in the human brain: evidence of age-related alterations

**DOI:** 10.1101/2025.10.12.681536

**Authors:** Kelly Perlman, Chuck T. Chen, Mackenzie E. Smith, John Kim, Gustavo Turecki, Richard P. Bazinet, Naguib Mechawar

## Abstract

The uncinate fasciculus (UF) is a long-range association fiber tract that serves to connect the anterior temporal lobe with the orbitofrontal cortex. The UF has been implicated via neuroimaging studies in the neurobiological vulnerability to psychiatric disorders posed by a history of childhood abuse (CA), as well as in the psychopathology underlying depressive disorders. Since the myelin sheath is highly enriched in lipids, white matter (WM) dysfunction may reflect alterations in the myelin lipid profile. In fact, our previous work showed that in the anterior cingulate cortex WM, there was a specific effect of CA in the choline glycerophospholipid fatty acids (FA) involved in the synthesis of arachidonic acid. Given that the UF does not exist in rodents, its molecular properties are highly understudied and its lipid composition is virtually unknown. As such, we sought to quantify the phospholipid FA and cholesterol quantities of the human postmortem UF and measure whether we could detect lipid-related or myelin-constituent gene/protein changes associated with CA and/or depression. Fresh-frozen left hemisphere UF samples were analyzed from individuals with depression who died by suicide with a history of severe CA (DS-CA), individuals with depression who died by suicide without a history of CA (DS), and non-psychiatric control subjects who died naturally or accidentally (CTRL). Phospholipids were separated by thin-layer chromatography. FA and non-derivatized cholesterol were quantified using gas chromatography-flame ionization detection. Relative expression of myelin-constituent genes (*PLP1*, *MAG*, *CNP*, *MOG*, *PLLP*, *MBP*, and *MOBP*) was measured by RT-qPCR, and levels of myelin-constituent proteins (MAG, MOG, MBP, and PLP) were measured by immunoblotting. We found no robust relationships between depression or CA and any lipid measures, nor in myelin-constituent gene and protein levels. However, in the phospholipids, we observed striking age relationships that varied across fractions, with an overall pattern of increases in monounsaturates and decreases in long chain omega-6 polyunsaturates with age. In tandem, we observed that most myelin-constituent genes and proteins showed decreasing trends with age, with *PLP1* and MAG showing significantly decreasing relationships. We hypothesize that the changes in lipid composition and lipid-protein interactions contribute to age-related myelin deficits and declines in cognition. The absence of group differences highlights the importance of regional specificity in molecular studies assessing neurobiological correlates of psychiatric disorders.

## Introduction

Approximately 5% of all human genes are devoted to lipid synthesis (van Meer et al., 2008) and when compared to non-neuronal tissues, the brain demonstrates a highly distinct lipid profile (Bozek et al., 2015). In human neocortex, concentrations of brain-enriched lipids have evolved three times faster when compared to chimpanzees (Bozek et al., 2015), pointing to an evolutionary significance for lipids in higher order cognitive function. White matter (WM) tracts, or fiber bundles, are collections of myelinated axons that transmit electric signals across distal brain regions and serve as “information highways”. The myelin sheath is comprised of 70-85% lipids according to its dry weight (Williams & Deber, 1993), and the specific composition and arrangement of lipids is known to influence its membrane properties and therefore its functional roles in brain circuits (Poitelon et al., 2020). For example, changes in myelin lipids such as the quantity of cholesterol and composition of fatty acids (FA) in phospholipids can impact its membrane properties, including compactness, stability, and permeability (Chrast et al., 2011; Naudí et al., 2015; Poitelon et al., 2020). Myelin lipids are estimated to be comprised roughly of 40% phospholipids, 40% cholesterol, and 20% glycolipids (O’Brien, 1965)

Despite their importance in both health and disease, there is very little characterization of long-range white matter tracts across the lifespan in the human brain. One particularly interesting and understudied WM tract is the uncinate fasciculus (UF), a long-range association fiber bundle that connects the anterior temporal lobe with the orbitofrontal cortex. Since this tract does not exist in rodents, its molecular properties have yet to be characterized. Furthermore, the development of the UF is highly protracted and it is one of the last tracts to fully mature with respect to its microstructural properties. Finally, it has been associated with both the underlying neurobiology of childhood trauma and the pathophysiology of depression via neuroimaging studies (Eluvathingal et al., 2006; Gur et al., 2019; Hanson et al., 2015; Ho et al., 2017; Xu et al., 2023). Therefore, a comprehensive characterization of this tract is warranted. Dysregulation in brain WM across a variety of limbic regions has been reported in both depression and in individuals with a history of childhood abuse (CA). However, it is unclear whether WM changes can be considered part of the pathophysiology of depression or whether they represent a biologically embedded risk factor for future psychopathology associated with experiencing CA. Neuroimaging studies that show depression-associated WM changes often fail to collect relevant information about childhood trauma. However, one study that investigated both depression and childhood trauma showed an interaction effect where individuals with severe depressive symptoms and childhood neglect had lower fractional anisotropy in the left UF (Tatham et al., 2016). Importantly, our previous work has shown the depressed suicides with a history of CA is associated with increased concentration of FAs in the arachidonic acid (ARA) synthesis pathway and corresponding gene expression changes in anterior cingulate cortex (ACC) gray matter (Perlman et al., 2021).

Therefore, this study had 2 complementary aims. Firstly, to perform a fundamental, in-depth characterization of phospholipid FA profiles and cholesterol composition of the UF. Secondly, with the addition of complementary measurements of key myelin-constituent genes and proteins, to examine if CA- or depression-related changes are observed in the UF.

## Methods

### Subject information

This research received ethical approval from the Douglas Mental Health University Institute’s Research Ethics Board. Group-matched brain samples, acquired with informed consent from the donors’ next of kin, were provided by the Douglas-Bell Canada Brain Bank. Fresh-frozen UF tissue was dissected from the left hemisphere. Data from the coroner’s office such as cause of death, time of death, and blood toxicology were obtained along with medical records when available. This information was complemented with data collected during a standard psychological autopsy procedure, as described elsewhere (Dumais et al., 2005; Perlman et al., 2021). Adapted versions of the Structured Clinical Interview for DSM-IV (SCID) and Childhood Experience of Care and Abuse (CECA) questionnaires were administered as part of this procedure. A panel of clinicians which includes a psychiatrist, synthesized all the evidence available to converge on an Axis I diagnosis, and which can be used to classify brain donors into 3 groups: depressed suicides with history of CA (DS-CA), depressed suicides without CA (DS), and non-psychiatric controls (CTRL). Neurodegenerative disorders constituted an exclusion criterion for all groups.

For all the gene and protein data, the UF was dissected at the point of intersection between the frontal lobe and the temporal lobe, lateral to the amygdala. Due to tissue availability limitations, all the lipid data was generated from the UF dissected at Brodmann area 38 (temporal pole).

These dissection sites were selected because the UF could be clearly distinguished from other concurrent tracts such as the inferior frontal occipital fasciculus. The samples were stored at −80 °C until biochemical analyses.

### Phospholipid FA quantification

For the phospholipids, we modified the protocol from our previous study (Perlman et al., 2021), to include all major phospholipid classes: choline glycerophospholipids (ChoGpl), ethanolamine glycerophospholipids (EtnGpl), phosphatidylserine (PtdSer), phosphatidylinositol (PtdIns), and sphingomyelin (CerPCho). Briefly, 50-70 mg of fresh-frozen UF tissue was dissected per sample. Adapted Folch method was used to extract total lipids (Folch et al., 1957), at which internal standard cocktail containing di-17:0 ChoGpl, di-17:0 EtnGpl, and 5-alpha cholestane was added (Supplementary Table 1). Then, 70 µL of the 500 µL total lipid extract (TL) was loaded into scored lanes of silica H-plates to perform thin layer chromatography (TLC). The TLC mobile phase was composed of 30:9:25:6:18 chloroform: methanol: 2-propanol: 0.25% KCL: triethylamine(v/v). Phospholipid bands were visualized under UV light after being sprayed with 0.1% (w/v) 8-anilino-1-naphthalene sulfonic acid and identified using a reference standard. Specific amounts of 17:0 free fatty acid was added as internal standards to the tubes containing PtdIns, PtdSer, and CerPCho (Supplementary Table 1) as purified form-specific internal standards were commercially unavailable. Each fraction was methylated with 14% BF_3_-MeOH, and all were incubated for 60 minutes, except for CerPCho which was incubated for 90 minutes, all at 100 °C. Total lipids (TL) was analysed via GC-FID as well. Samples were measured by gas chromatography-flame ionization detection (GC-FID) (Varian 430 gas chromatograph (Bruker, Billerica, MA, USA). Fatty acid methyl esters were eluted on a DB-FFAP column (30m x0.25 mm i.d. x 0.25μm film thickness) (J&W Scientific, Agilent Technologies). Run program was set at initial temperature of 50°C (1 min), 130°C ramp at 30°C/min, 175°C ramp at 10°C/min, 230°C ramp at 5°C/min (9.5 min hold), and final 240°C at 5oC/min (11.13 min hold). The TL, ChoGpl, and EtnGpl fractions were run on split injection mode, while CerPCho, PtdSer, and PtdIns fractions were run on splitless injection mode.

Chromatograms were analyzed with the CompassCDS software and annotated based on the GLC 569 external reference standard. Concentrations of each FA in µg/g were calculated based on peak comparison to internal standard. Furthermore, relative percentages of each FA were calculated. Peaks that could not be reliably annotated were excluded – for example, C20:5n-3 and C20:2n-6 were detected at trace amounts in CerPCho but not reliably enough to quantify.

### Cholesterol Quantification

Non-derivatized cholesterol were run on a HP-5 ms capillary column (30 m x 0.25 mm i.d. x 0.25 μm d_f_) (J&W Scientific, Agilent Technologies) using a Varian 430 GC-FID. The injector and detector ports were set to 250°C and 300°C, respectively. Cholesterol was eluted using a temperature program set initially at 100°C for 1 min, followed by increases at 15°C/min for 17 min until reaching final temperature of 280°C and complete the run of 30 min. The carrier gas was helium set at a constant flow rate of 1 ml/min. Cholesterol was quantified using peak comparison to the internal standard, 5α-cholestane. Annotations were performed on CompassCDS.

### Ratios and indices

We also computed several relevant ratios and indices, including the ratio of omega-6/omega-3 FAs and the quantity of highly unsaturated FAs (HUFA; sum of FAs with 20 or more carbons and 3 or more double bonds). Then, we calculated a verity of indices, including the peroxidation index (Hulbert et al., 2007), which quantifies the susceptibility of a given lipid to oxidation, with the following formula:

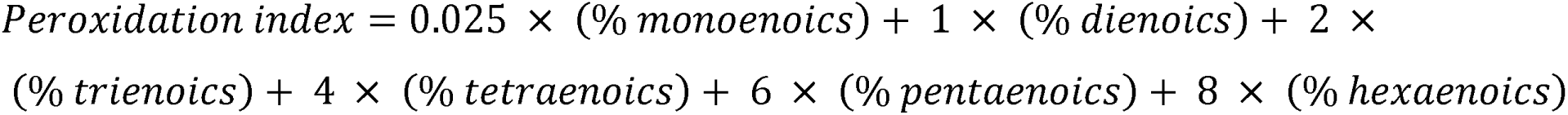

Furthermore, we calculated the unsaturation index (Hulbert et al., 2007) with the following formula:

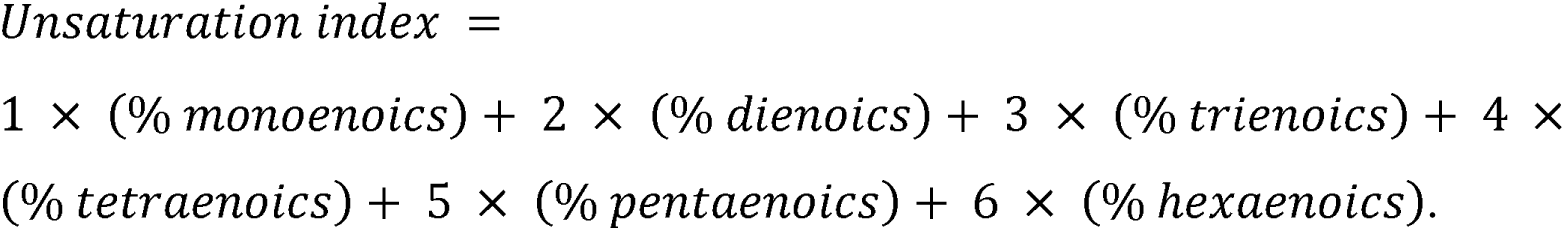

The final index calculated was the chain length index with the formula:

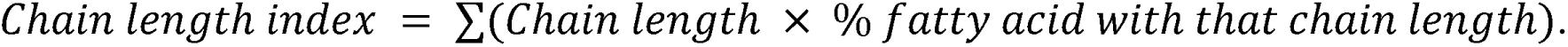

### Expression of myelin-constituent genes

RNA was extracted from homogenized UF tissue in Qiazol lysis buffer using the Qiagen RNEasy Lipid Tissue Mini Kit. DNA was removed with Qiagen RNase-Free DNAse to the extracted RNA. A standardized amount of purified RNA from each UF sample was reverse transcribed with MLV Reverse Transcriptase Kit yielding UF cDNA for each subject. A standard curve was made to determine the relative amount of cDNA amplification. The forward and reverse primers were diluted and mixed with Applied Biosystems PowerUp SYBR Green master mix. RT-qPCR was carried out with the Applied Biosystems QuantStudio 6 Flex PCR machine. The output was analyzed with the QuantStudio Real-Time PCR software. Replicates with an expression higher than ± 0.3 standard deviation from the mean expression values were withdrawn from analysis. See Supplementary Table 2 for details of the primers used in RT-qPCR.

### Expression of myelin-constituent proteins

Protein lysates for each subject were generated by homogenizing UF tissue in RIPA buffer. Protein concentration of each lysate was measured using the BCA Protein Assay Kit (Thermofisher) and concentration was determined with a Tecan Spark10M plate reader. An equal quantity of protein for each subject was run on a BioRad Mini-PROTEAN TGX Stain-Free 4-20% gel and separated by way of SDS-PAGE. The electrophoresis was run at 135V for approximately 60 minutes at ambient temperature. Next, the proteins in the gel were transferred onto an Amersham Protran Premium 0.2 μm nitrocellulose blotting membrane with the BioRad Trans-Blot Turbo Transfer System. This membrane was imaged in the BioRad Chemidoc Touch Imaging System to obtain a quantification of total protein expression.

The membrane was blocked for 1 hour with 5% skim milk, then primary antibodies in 1% skim milk blocking solution were added and incubated overnight at 4 °C. Then, HRP-conjugated secondary antibodies in 1% skim milk were incubated for 1 hour at room temperature. The antibody details are listed in Supplementary Table 3. After the incubations, the membranes were exposed to electrochemiluminescent (ECL) fluid for 2 minutes, and the chemilumiscent signal generated was captured with the BioRad Chemidoc Touch Imaging System imager and measured using the ImageLab software.

### Statistics

Statistics were performed using R version 4.4.2. ANCOVA models were constructed with the following factors: group, age, sex, brain pH, and postmortem interval (PMI), and the presence/absence of a documented lipid-related condition. For this study, specific attention was made to the donor’s documented lipid-related conditions including any listed of elevated cholesterol, coronary artery disease, dyslipidemia, or type 2 diabetes or medications to treat those conditions. This is relevant, as it has been shown that statins lower brain cholesterol by way of reducing both *de novo* synthesis and turnover rate (Cibičková, 2011). If the donor either had an antidepressant prescription in 3 months before their death or if they had antidepressants detected in their body at time of death, they were considered to be “positive” for antidepressants. P-values were corrected for multiple comparison within each predictor across FAs within in a phospholipid fraction using the Benjamini-Hochberg (BH) method.

Based on the literature, we hypothesized that age relationships in lipids are often non-linear, and so for the age-specific analyses, a multi-step approach was used. First, regressions were fitted for linear, quadratic, cubic, and logarithm relationships. Then, with the approach in (Perlman et al., 2025), whichever model minimizes the Bayesian information criteria (BIC) was considered to be the best fitting. However, since a BIC value difference less than 2 does not inform the selection of one model over another (Berchtold, 2010), we selected the linear model as the “default” due to its interpretability, and only chose a non-linear model when if BIC_linear_ – BIC_other_ ≥ 2. P < 0.05 was selected as the threshold for nominal statistical significance across analyses. Pearson’s correlations were performed and represented in correlation plots (R corrplot package). Due to the exploratory nature of the correlation analyses, correlation test p-values were not corrected for multiple comparisons. Data in the figures are represented as mean ± standard error of the mean.

## Results

### Fundamental characterization of UF lipids

Supplementary Table 4 contains the subject information for the UF lipids cohort (n=80). The mean total concentration for each of the 5 main phospholipid fractions, cholesterol, and the TL are plotted in Figure 1A. Of the phospholipids, EtnGpl was the most abundant followed closely by ChoGpl while PtdIns was the least abundant phospholipid. The mean cholesterol concentration was 12611.59 ± 5989.21 µg/g, which is the most abundant individual lipid species measured. The distribution of FA classes of each phospholipid and the TL are illustrated in Figure 1B. The FA class distribution in each isolated phospholipid fraction matches the expected overall pattern based on the literature of WM (e.g., EtnGpl is enriched in omega-6 FA and CerPCho is enriched in saturated FA/SFA). The concentration data for each FA for each phospholipid fraction as well as the TL (represented as mean ± standard deviation) is available in Table 1. In the TL, the most abundant FA is C18:1n-9 (OLA, 8468.14 ± 1925.87 µg/g) followed by C18:0 (STA, 6086.21 ± 821.10 µg/g), then C16:0 (PAM, 4974.11 ± 608.19 µg/g), C22:4n-6 (AdA, 2891.96 ± 925.67 µg/g), and C20:4n-6 (ARA, 2209.72 ± 495.80 µg/g). When performing a principal component analysis on the FAs detected in every phospholipid fraction, the fractions clearly separate along the first 2 principal components (Supplementary Figure 1), reflective of their distinct FA “signatures”.

**Figure 1.**
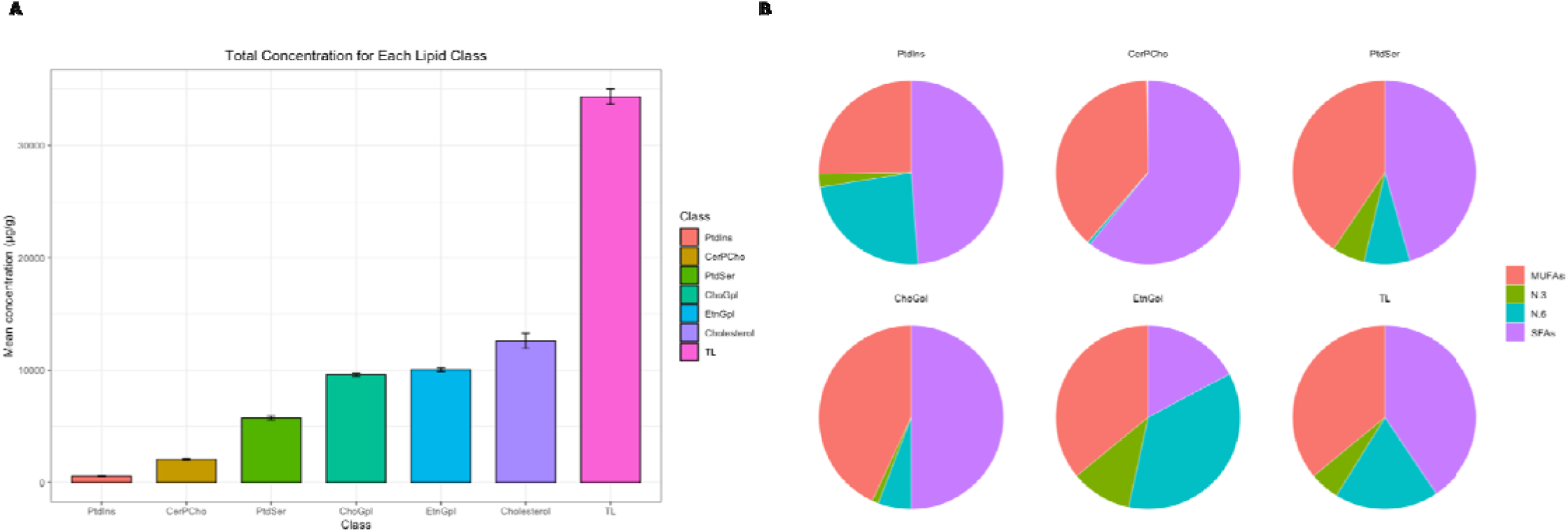
Mean totals and fraction-specific characterization of FA classes. **A)** Bar plot showing the mean concentration ± standard error for the totals of all measured phospholipid fractions, cholesterol, and the TL in ascending order. TL refers to the total FA, not just the FAs derived from phospholipids. Therefore, the FAs in the TL also come from glycolipids, diacylglycerols, and other lipid species. **B)** Pie charts showing the mean relative percentage of MUFA (pink), omega-6 (teal), omega-3 (green), and SFA (purple) FAs for each phospholipid fraction as well as the TL. MUFA: monounsaturated fatty acid, SFA: saturated fatty acid. PtdIns: phosphatidylinositol, PtdSer: phosphatidylserine, CerPCho: sphingomyelin, ChoGpl: choline glycerophospholipids, EtnGpl: ethanolamine glycerophospholipids. TL: total lipid.

**Table 1:**

| Fatty acid | ChoGpl | EtnGpl | PtdIns | PtdSer | CerPCho | TL |
| --- | --- | --- | --- | --- | --- | --- |
| C 14:0 | 84.45 ± 14.19 | 13.37 ± 5.10 | 2.08 ± 1.55 | 3.16 ± 1.28 | 4.62 ± 1.90 | 98.38 ± 26.68 |
| C 16:0; PAM | 3491.83 ± 422.61 | 445.39 ± 71.65 | 95.88 ± 47.76 | 102.67 ± 32.28 | 129.10 ± 43.44 | 4974.11 ± 608.19 |
| C 18:0; STA | 1191.04 ± 163.53 | 1218.26 ± 356.44 | 166.84 ± 102.84 | 2491.91 ± 774.84 | 805.75 ± 247.94 | 6086.21 ± 821.10 |
| C 20:0 | 7.69 ± 1.61 | 15.15 ± 4.37 | 1.05 ± 0.54 | 10.20 ± 2.60 | 33.90 ± 10.23 | 101.72 ± 36.97 |
| C 22:0 | 2.34 ± 0.76 | 5.30 ± 1.54 | 0.81 ± 0.33 | 4.15 ± 1.20 | 41.16 ± 15.97 | 96.63 ± 67.19 |
| C 23:0 | ND | ND | 0.61 ± 0.32 | ND | 54.08 ± 22.10 | 127.93 ± 65.78 |
| C 24:0 | 6.90 ± 4.46 | 8.24 ± 5.62 | 2.16 ± 1.02 | 8.58 ± 34.39 ** | 164.58 ± 73.57 | 2194.44 ± 827.68 |
| Total SFAs | 4784.24 ± 559.92 | 1705.71 ± 370.04 | 269.43 ± 142.71 | 2620.68 ± 798.89 | 1233.18 ± 373.98 | 13729.69 ± 1725.42 |
| C 16:1n-7 | 119.26 ± 25.55 | 44.55 ± 19.78 | 4.02 ± 2.04 | 7.75 ± 2.50 | 1.33 ± 0.52 | 1119.12 ± 402.35 |
| C 18:1n-7 | 617.46 ± 101.66 | 449.11 ± 120.35 | 36.62 ± 19.71 | 222.51 ± 70.62 | 4.66 ± 1.77 | 1562.66 ± 325.21 |
| C 18:1n-9 (OLA) | 3327.61 ± 608.87 | 2678.00 ± 778.62 | 82.60 ± 50.78 | 1918.89 ± 653.95 | 27.30 ± 7.96 | 8468.14 ± 1925.87 |
| C 20:1n-9 | 67.07 ± 17.13 | 435.52 ± 157.47 | 5.98 ± 4.24 | 200.28 ± 79.58 | 1.61 ± 0.64 | 762.52 ± 285.36 |
| C 22:1n-9 | 10.28 ± 4.40 | 37.65 ± 13.24 | 5.27 ± 3.35 | 13.17 ± 4.57 | 9.37 ± 9.07 | 71.86 ± 29.88 |
| C 24:1n-9 | 10.64 ± 3.78 | 6.92 ± 2.50 | 0.98 ± 0.63 | 4.79 ± 1.68 | 767.18 ± 309.15 | 229.46 ± 111.62 |
| Total MUFAs | 4152.32 ± 731.78 | 3651.76 ± 1058.77 | 135.46 ± 75.62 | 2367.38 ± 796.84 | 811.43 ± 314.72 | 12433.13 ± 2947.73 |
| C 18:2n-6; LNA | 65.58 ± 15.36 | 24.88 ± 6.12 | 4.24 ± 2.57 | 6.84 ± 2.28 | 1.97 ± 0.51 | 125.87 ± 29.38 |
| C 20:2n-6 | 8.87 ± 9.45 | 24.69 ± 11.53 | 1.37 ± 1.23 | 7.02 ± 3.15 | ND | 161.07 ± 45.94 |
| C 20:3n-6; DGLA | 49.09 ± 11.60 | 134.87 ± 38.02 | 12.55 ± 9.49 | 40.34 ± 14.81 | 5.49 ± 2.11 *<br>co-eluted with C21:0 | 312.20 ± 86.98 |
| C 20:4n-6; ARA | 341.26 ± 69.69 | 1257.35 ± 215.24 | 109.15 ± 79.98 | 122.63 ± 37.30 | 2.64 ± 1.54 | 2209.72 ± 495.80 |
| C 22:2n-6; DDA | 6.68 ± 2.87 | 19.90 ± 7.14 | 1.08 ± 0.47 | ND | 1.81 ± 1.55 | 375.41 ± 171.98 |
| C 22:4n-6; AdA | 73.79 ± 11.30 | 2053.30 ± 433.09 | 18.47 ± 14.45 | 230.68 ± 65.10 | 3.53 ± 1.68 | 2891.96 ± 925.67 |
| C 22:5n-6; DPA n-6 | 10.10 ± 3.60 | 117.70 ± 41.52 | 1.18 ± 0.88 | 42.68 ± 20.14 | 0.40 ± 0.20 | 233.44 ± 87.76 |
| Total N-6 | 555.38 ± | 3632.68 ± | 148.03 ± | 450.21 ± | 10.35 ± | 6310.38 ± |
|  | 95.31 | 572.61 | 103.75 | 125.32 | 3.68 | 1570.11 |
| C 20:5n-3;<br>EPA | ND | ND | ND | ND | ND | ND |
| C 22:5n-3;<br>DPA n-3 | 5.78 ±<br>1.61 | 63.40 ±<br>18.04 | 0.72 ±<br>0.54 | 9.61 ±<br>2.88 | ND | 127.81 ±<br>54.15 |
| C 22:6n-3;<br>DHA | 97.76 ±<br>26.34 | 985.89 ±<br>381.64 | 11.86 ±<br>7.39 | 284.23 ±<br>142.19 | ND** | 1723.21 ±<br>828.39 |
| Total N-3 | 103.54 ±<br>27.19 | 1049.28 ±<br>385.77 | 12.58 ±<br>7.88 | 293.84 ±<br>143.59 | ND** | 1851.02 ±<br>855.90 |
| Total FA | 9595.48 ±<br>1243.32 | 10039.43 ±<br>1434.06 | 565.51 ±<br>319.92 | 5732.12 ±<br>1665.76 | 2060.45 ±<br>663.03 | 34324.21 ±<br>6025.21 |

Concentration (µg/g) data is represented as mean ± standard deviation. ND: non-detectable.
\*co-elution with C21:0
\*\* co-elution with C24:0 (~1% of total represents DHA). As such, omega-3 quantities in CerPCho are negligible.

We specifically validated the peak of C24:0 in CerPCho with GC-MS (as described in (Lacombe et al., 2023; Metherel et al., 2018)) with a specific target ion for C22:6n-3 [DHA]) (Supplementary Figure 2) because the retention time matched closest to DHA on the 569 external reference standard. From running six representative samples, we found that these peaks co-elute and, on average, 98.9% of the total area matched to C24:0, and 1.1 % to C22:6n-3, with these patterns highly consistent across all individuals in the CerPCho fraction (Supplementary Table 2). We therefore annotated the peak as C24:0, but it is worth noting that there is in fact a small amount of DHA in SM. If the mean concentration of that peak is 164.58 µg/g, then 1.1 % of that would be 1.81 µg/g which is an estimate to the amount of DHA in CerPCho of the UF.

Furthermore, we observe that the summary ratios and indices (e.g.,chain length index, unsaturated index, peroxidation index, omega-6/omega-3 ratio, and HUFA) correspond with the known FA makeup of these phospholipids. For example, CerPCho shows the highest chain length index and lowest unsaturated index of all the fractions, consistent with CerPCho being enriched with long FA tails, which are primarily saturated (Slotte, 2016). The average values for the ratios and indices for each phospholipid fraction and the TL, can be found in Supplementary Table 7.

### Group differences in characterization of UF lipids

The distribution of phospholipid fractions between groups (i.e., the ratio of total FAs per fraction) are demonstrated in Figure 2A. With respect to the phospholipids, several FA showed statistically significant group differences but did not retain significance upon correction for multiple comparisons (Supplementary Figure 3). Specifically, group was nominally significant for PtdSer C20:3n-6 relative percentage (p = 0.019) and the TL C20:3n-6 relative percentage (p = 0.0087). Importantly, the changes were specific to the DS group (not both DS and DS-CA), and for FA of very small quantities (less than 1%), thus the biological relevance of these findings is unknown. We found no other changes in FA quantities (either concentration or relative percentage) associated with CA or with depression.

**Figure 2.**
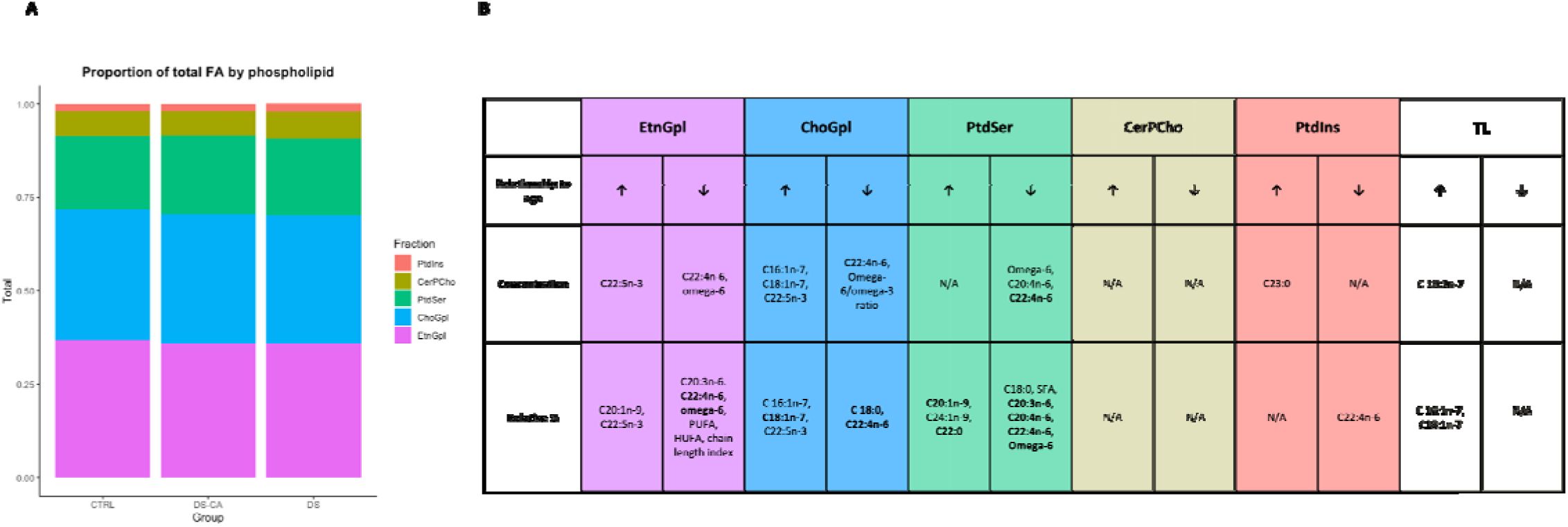
UF phospholipids show no robust effect of group but striking effect of age. **A**) Stacked bar charts show the proportion of the total of each phospholipid class by group, as measured by total FA for each class. **B)** Summary table showing age related changes in lipids separated by phospholipid fraction and TL. Arrows pointing upward indicate a positive coefficient for age, and arrows pointing downwards indicate a negative coefficient for age. FAs or summary metrics in bold indicate statistical significance after correction for multiple comparisons (BH-corrected p < 0.05).

We did not find any differences between groups in cholesterol concentration (CTRL: 12242.92 ± 804.48 µg/g; DS: 12378.99 ± 1204.62 µg/g; DS-CA: 13114.49 ± 1229.06 µg/g). However, several studies have observed that individuals who died by suicide using violent methods have lower serum cholesterol than those who died by suicide using non-violent methods (Aguglia et al., 2019; Alvarez et al., 2000; Golomb, 1998), and one study found that gray matter cholesterol in the frontal cortex is lower in postmortem brains of depressed individuals who died by way of violent suicide as compared to depressed suicides who died by non-violent methods and controls (Lalovic et al., 2007). When stratifying by violent vs. non-violent method of suicide, we still did not identify group differences. Therefore, cholesterol concentration in the UF is not associated with CA or depression.

### Age relationships

While we did not observe much of a relationship between FA quantities and group, we did observe a series of striking patterns between FA quantities and age, spanning a wide age range (15-85). The age relationships are summarized in Figure 2B. PtdSer and EtnGpl showed the most age-related changes, while CerPCho showed none. For ChoGpl, directly proportional relationships were observed for C16:1n-7 (concentration p=0.039, relative percentage p=0.0094), C18:1n-7 (concentration p=0.0468, relative percentage p=0.0021), C22:5n-3 (concentration p=0.0122, relative percentage p=0.0148), and inversely proportional relationships were observed for C18:0 (relative percentage p = 0.0008), C22:4n-6 (concentration p=0.024, relative percentage p=0.0004), omega-6/omega-3 ratio (concentration p = 0.0091), summarized in Supplementary Figure 4. For EtnGpl, directly proportional relationships with age were observed for C20:1n-9 (relative percentage p=0.043), C22:5n-3 (concentration p= 0.0496, relative percentage p =0.032), and inversely proportional relationships were observed for C20:3n-6 (relative percentage p =0.045), C22:4n-6 (concentration p =0.049, relative percentage p=0.0009), omega-6 (concentration p =0.040, relative percentage p<0.0001), PUFA (relative percentage p =0.011), HUFA (relative percentage p=0.012), chain length index (relative percentage p =0.0067), summarized in Supplementary Figure 5. For PtdSer, directly proportional relationships with age were observed for C20:1n-9 (relative percentage p=0.0065), C24:1n-9 (relative percentage p=0.045), C22:0 (relative percentage p=0.00001) and inversely proportional relationships were observed for C18:0 (relative percentage p=0.024), SFA (relative percentage p=0.025), C20:3n-6 (relative percentage p=0.0005), C20:4n-6 (concentration p=0.038, relative percentage p<0.00001), C22:4n-6 (concentration p=0.0058, relative percentage p<0.00001), omega-6 (concentration p=0.018, relative percentage p=0.0004), summarized in Supplementary Figure 6. For PtdIns, a directly proportional relationship with age was observed for C23:0 (concentration p=0.031) and inversely proportional relationship was observed for C22:4n-6 (relative percentage p=0.021), summarized in Supplementary Figure 7. No age-related changes were observed for any FA in CerPCho (either concentration or relative percentage). For the TL, directly proportional relationships were observed for C16:1n-7 (concentration p=0.015, relative percentage p=0.0049) and C18:1n-7 (relative percentage p=0.040), summarized in Supplementary Figure 8. The age term for C18:0 ChoGpl concentration was best fit by a logarithmic model. All the other significant age terms were best fit by linear models. No significant age relationships were observed for cholesterol. The FAs that survived correction for multiple comparison (BH corrected p < 0.05) are bolded in Figure 2B.

### Regional comparison of ChoGpl

We previously published data from the ChoGpl fraction of the human ACC white matter (Perlman et al., 2021). Thus, to understand whether we observe differences in the same phospholipid fraction across brain regions, we compared the concentrations of the ACC and UF ChoGpl fractions (Supplementary Table 8) of the overlapping FAs in overlapping subjects between the two studies (n=70). All FAs, except certain trace FAs (<1% of total FA) overlapped. The correlation between the average ACC and UF ChoGpl FA concentrations was nearly perfect (r = 0.993). Despite this near-perfect correlation, the two regions still show clear separation across a PCA plot (Supplementary Figure 9). While batch effects cannot be ruled out, there are differences in the FA composition that can explain the separation. For example, in the UF, the concentrations of PAM (C16:0, 3490.12 ± 424.73) and OLA (C18:1n-9, 3400.93 ± 578.75) are roughly equal, but in the ACC, there is more OLA (4066.80 ± 690.47) than PAM (3397.46 ± 614.38). Furthermore, in the UF, DHA is the 7^th^ most abundant FA, while in the ACC, DHA is the 10^th^ most abundant. When comparing the age patterns of ChoGpl FAs across both regions, we observe both similarities and differences. In ChoGpl in ACC WM, total FA concentration is decreasing with age in addition to most other FAs, with no FAs showing increasing concentrations (Supplementary Figure 10). In UF, the changes observed in this subset of the cohort (n=70) are similar to those observed in the full cohort (excluding those for C22:5n-3 and omega-6/omega-3 ratio which did not quite reach significance) and are not reflective of overall lower ChoGpl content. In other words, the changes in FA concentration with age in ACC are indicative of a more global effect (i.e. significant decreases in most FAs), while the changes in the UF concentration are more specific (C16:1n-7, C18:1n-7 increasing, decrease in AdA). Across both regions, we observed striking significant AdA decreases in concentration and relative percentage metrics with age, and significant increase in C16:1n-7 and C18:1n-7 relative percentage with age. It is worth noting that AdA was the FA most strongly associated with aging in the ACC ChoGpl by absolute correlation coefficient (Perlman et al., 2021), and AdA also decreases in UF ChoGpl, in addition to UF EtnGpl, PtdSer, and PtdIns. Thus, AdA might be more heavily influenced by the aging process across brain regions.

### Myelin-constituent genes and proteins

Just as the particular lipid composition of myelin is crucial to its function, so too are the specific proteins that make up the sheath. As such, we sought to examine several myelin-constituent proteins via western blot (MAG, MBP, MOG, PLP) and the genes that code for them via RT-qPCR (CNP, MAG, MBP, MOBP, MOG, PLLP, PLP1). The subject information for qPCR cohort can be found in Supplementary Table 5 and the western blot subject information can be found in Supplementary Table 6 though these cohorts are largely overlapping. No statistically significant differences between groups were observed for any of the genes (Figure 3A) or proteins (Figure 3B) measured, indicating no robust association with depression or CA. With respect to age, PLP1 gene expression was significantly associated with age (p = 0.032), though this did not retain significance after correction for multiple comparisons (p = 0.081). Nevertheless, all genes measured (except MOBP) show a decrease across the age span (Figure 3C). Of the proteins measured, MAG was significantly associated with age (p = 0.022), though this did not retain significance after correction for multiple comparisons (p = 0.11). Descriptively, however, MAG, MBP, and MOG show decreases across the age span, while PLP protein appears steady (Figure 3D). Overall, the trend of myelin constituent-gene and protein expression is a decline with age.

**Figure 3.**
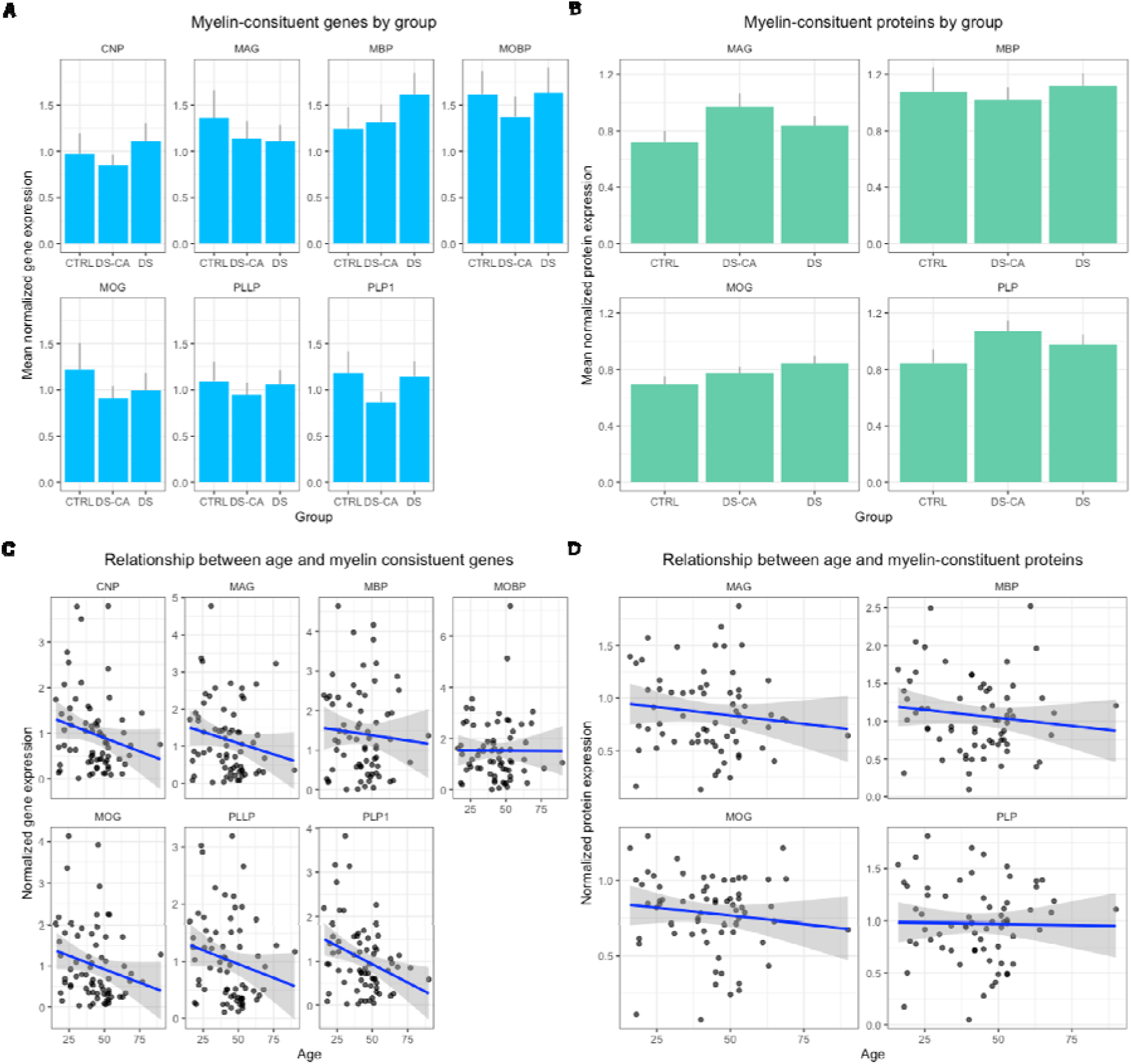
Myelin-constituent genes and proteins differ by age, not by group. **A**) Bar plots showing mean + standard error of gene expression values (normalized to GAPDH) for CNP, MAG, MBP, MOBP, MOG, PLLP, and PLP1 across all 3 groups with no statistically significant differences. **B**) Bar plots showing mean + standard error of protein expression values (normalized to total protein quantity) for MAG, MBP, MOBP, MOG, and PLP across all 3 groups no statistically significant differences. **C**) Scatter plot with trend line (blue) showing age plotted against gene expression values (normalized to GAPDH), with PLP1 showing a significant negative relationship (p = 0.032). **D**) Scatter plot with trend line (blue) showing age plotted against protein expression values (normalized to total protein quantity), with MAG showing a significant negative relationship (p = 0.022). CTRL: control, DS: depressed suicides, DS-CA: depressed suicides with a history of CA, GAPDH: glyceraldehyde 3-phosphate dehydrogenase, CNP: 2’,3’-cyclic nucleotide 3’-phosphodiesterase, MAG: myelin-associated glycoprotein, MBP: myelin basic protein, MOBP: myelin-associated oligodendrocyte basic protein, MOG: myelin oligodendrocyte glycoprotein, PLLP: plasmolipin, PLP1: proteolipid protein.

We then examined the correlations between summary lipid metrics (e.g. SFAs, MUFAs, etc.) and gene or protein expression for subjects that were overlapping between the datasets (n=44).

Broadly, the quantities of all lipids are positively correlated with all myelin protein expression. If a greater concentration of lipids is indicative of more myelin overall, then it follows that more protein would be required as well to maintain the proper stoichiometric relationship (Figure 4). This broadly positive correlation does not hold when assessing lipid concentration and gene expression, where there is a mix of positive and negative correlations. In the relative percentage metric, MUFA percentage was largely negatively correlated with myelin genes, while SFA percentage was largely positively correlated (Supplementary Figure 11). The most striking observation is that the myelin proteins, but not the genes, appear to be strongly correlated with PtdIns quantities both in concentration and relative percentage, as seen by the consistent statistical significance and strong correlation coefficients. Interestingly, in relative percentage metrics, PtdIns MUFA are significantly negatively correlated with myelin proteins, in contrast to all other PtdIns lipid classes which are significantly positively correlated. This strong myelin protein-PtdIns correlation is possibly due to the key signalling role of phosphorylated PtdIns in trafficking myelin proteins to their proper location in the sheath (Baron et al., 2015; Baskin et al., 2016; De Craene et al., 2017; Mironova et al., 2016; Nawaz et al., 2009; Snaidero et al., 2014).

**Figure 4.**
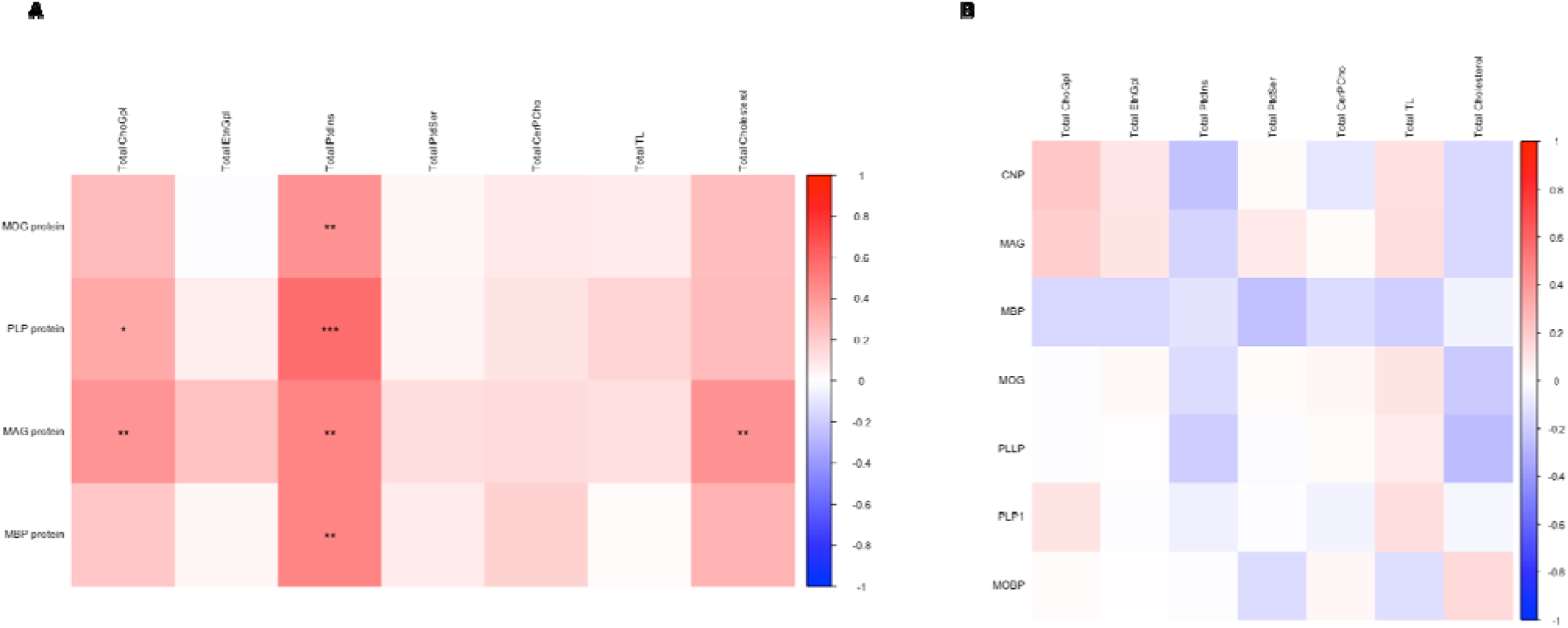
Correlations between total lipids and myelin-constituent proteins/genes. Correlation plots showing the relationships between total concentration (μg/g) of each lipid class (phospholipid fractions, the TL, and cholesterol) and myelin-constituent genes **(A)** and proteins **(B).** Due to the exploratory nature of this analysis, p-values were not corrected for multiple comparisons. The color bar represents Pearson’s correlation coefficient between −1 and 1, where negative coefficients are blue in color and positive coefficients are red in color. ***p < 0.001, ** p < 0.01,* p <0.05.

## Discussion

In this study, we provide, to the best of our knowledge, the first characterization of the UF phospholipid FA and cholesterol profile, paired with measures of myelin-constituent genes and proteins, to examine relationships with CA, depression, and age. The importance of region-specific lipid characterization should not be overlooked. A lipidomic study of 75 regions of the adult human brain recently showed that the composition of 93% of the lipids measured were variable across brain regions (Osetrova et al., 2024), lend further support to the notion of regional specificity (Naudí et al., 2015). A recent spatial lipidomic study even described differences in lipid composition across WM tracts (Zavolskova et al., 2025). There is a paucity of human postmortem brain datasets for most WM fiber bundles, and the current characterization of UF lipid composition and comparison of the UF ChoGpl and ACC WM ChoGpl should constitute a useful resource to the field.

The absence of robust significant changes of UF FA or cholesterol in depression or CA concurs with the absence of changes found in myelin-consistent gene expression or protein levels. Taken together, it appears that the myelin in this region does not display any evident long-lasting impacts of depression or CA. These findings indicate that CA-related changes, at least, may be region-specific. In the ACC WM, we observed DS-CA group-specific increases of ChoGpl FAs that are part of the ARA synthesis pathway, and corresponding gene expression changes in ARA metabolism genes (Perlman et al., 2021). Furthermore, in the ACC, myelin-constituent genes were heavily downregulated, painting an overall picture of myelin dysregulation in this region (Lutz et al., 2017). As such, one should not generalize neurobiological findings of any given limbic areas in psychiatry, as they are likely to be region-specific, and likely circuit-specific. We speculate that the UF neuroimaging findings of CA might represent transient, dynamic changes in maturation rate, rather than long-lasting neurobiological correlates that can be observed postmortem, especially given that most studies are conducted in adolescents and young adults (Eluvathingal et al., 2006; Granger et al., 2021; Gur et al., 2019; Ho et al., 2017).

While each phospholipid fraction had its own aging “signature”, we observed a pattern in which MUFA (especially C16:1n-7 and C18:1n-7) increased with age and long chain omega-6 FA (especially AdA) decreased with age. In contrast, the UF had a remarkably steady concentration of cholesterol across a wide age range. A previous study of the human postmortem hippocampus suggested that while the de novo cholesterol synthesis rate may decrease upon aging, the absolute cholesterol content remains stays stable (perhaps due to a reduction in 24S-hydroxycholesterol levels, limiting cholesterol efflux from the brain) (Thelen et al., 2006). This stability of cholesterol content has been replicated in the corpus callosum of rhesus monkeys (Dimovasili et al., 2024). Though age-related decreases in brain cholesterol have been reported as well (Stommel et al., 1989), it is likely that cholesterol relationships with age are region specific (Söderberg et al., 1990).

Previous aging studies of both animal and postmortem human tissue are in general agreement with the FA results presented here (Carver et al., 2001; Furber et al., 2022; Mota-Martorell et al., 2022; Söderberg et al., 1990). A study of the orbitofrontal cortex (which sends and receives projections via the UF) found that C20:4n-6 decreases with age, and a study of the frontal cortex shows that both ARA and AdA (along with many other PUFAs) also decrease with age (Carver et al., 2001). Interestingly, our findings recapitulate those of the anterior corpus callosum in aging mouse brain as investigated with a completely different method, Fourier transform infrared spectroscopic imaging (Furber et al., 2022). These authors found that age was associated with an increase in MUFA in tandem with a decrease in PUFA, which parallels our results (Furber et al., 2022). It is interesting to consider the notion that the overall pattern of aging in FA classes may be conserved across species.

While most FA supplementation studies focus on omega-3s, there is some evidence that omega-6 supplementation may be beneficial in aging. In a double-blind trial of ARA supplementation among 25 elderly men, it was observed that 1 month of ARA supplementation (as compared to olive oil placebo) decreased P300 event related potential (ERP) latency and increased P300 ERP amplitude, indicating improved cognition (Ishikura et al., 2009). Studies in rats have shown that 1) aged rats have lower membrane ARA concentration, 2) lower ARA concentration is associated with lower hippocampal long-term potentiation, and 3) supplementing with ARA can reverse these hippocampal long term potentiation deficits and improve cognitive abilities (Bethlehem et al., 2022; Kotani et al., 2003; McGahon et al., 1997; Okaichi et al., 2005). Together, the evidence suggests that further research on omega-6 supplementation in aging is warranted.

Animal studies support our finding that myelin-constituent genes and proteins tend to decline with age (Xie et al., 2013; Ximerakis et al., 2019). The biophysical and biochemical interactions between lipids and proteins in the myelin sheath are absolutely essential to the specialized molecular organization of the membrane (Ozgen et al., 2016). For example, microdomains such as lipid rafts serve as platforms for myelin protein trafficking and hubs for cell signalling (Dupree & Pomicter, 2010). Myelin proteins interact with the myelin lipids to create and maintain structural integrity of the sheath (e.g., PLP anchors the external leaflets, MBP compacting the cytoplasmic leaflets, and MAG works as an adhesion molecule between the axon-myelin interface by binding with gangliosides) (Ozgen et al., 2016; Pronker et al., 2016). There are many complex lipid-protein interactions that are key to myelin biosynthesis, maintenance, and integrity over the long term (Barnes-Vélez et al., 2023; Ozgen et al., 2014, 2016). Correspondingly, our findings here show that several lipids, especially the signalling lipid PtdIns, are correlated with myelin-constituent proteins. Age-related declines in myelin integrity have been associated with age-related declines in cognition in the human brain (Coelho et al., 2021; Mendez Colmenares et al., 2024), including in the UF proper (Zahr et al., 2009). Thus, we speculate that age-related changes in myelin lipid and protein levels, as observed in this study, might impact lipid-lipid and lipid-protein interactions such that microstructural integrity of the myelin is impaired sufficiently to affect information transfer along axons.

It is worth noting the limitations of this study. Firstly, due to the low number of females in our cohort, consistent with the higher rates of suicide in males, we were likely unable to reliably detect sex differences (Turecki et al., 2019). Secondly, for the annotation of FAs, we only considered FA that were present on the GLC569 external standard (with the exception of C24:1n-9), suggesting that we could be missing changes in FAs that we did not consider (e.g., longer FAs). Third, we do not separate the ether phospholipids from the ester phospholipids. It is estimated that over 80% of EtnGpl in myelin is in plasmalogen form, thus we can assume that most of the EtnGpl measured here is ethanolamine plasmalogen (PlsEtn) (Barnes-Vélez et al., 2023). It seems as though choline plasmalogen are enriched in WM fractions as well, though less so than PlsEtn (Osetrova et al., 2024). Finally, we cannot ascertain whether alterations in FA are due to changes in synthesis, transport, turnover, oxidation, and/or dietary intake. We do not have information on diet or other lifestyle factors that could influence the brain lipidome, particularly PUFAs which are taken up from the periphery (Bazinet & Layé, 2014). In the future, it would be interesting to probe the dietary origins of PUFAs using natural abundance carbon isotope ratio analysis (^13^Carbon /^12^Carbon ratio) (Lacombe et al., 2023).

A key future direction would be to probe the glycolipid composition of the UF and other WM tracts. Glycolipids, especially the cerebroside galactosylceramidase and its sulfated form (sulfatide), are known to be highly enriched in the myelin sheath, and to play a key role in long-term myelin and axonal stability (Barnes-Vélez et al., 2023; Marcus et al., 2006). It would also be tremendously valuable to have both central and peripheral (e.g., serum, plasma) lipid measurements to assess to what degree the blood might reflect lipid changes in the brain and whether such peripheral measures could serve as biomarkers of health and disease.

Newer methods for quantification (e.g., flow cytometry assisted single-cell lipidomics (Hancock et al., 2023)) and visualization (e.g., MALDI-imaging mass spectrometry (Sugiura et al., 2009)) of lipids in individual cells will provide better resolution to answer questions regarding cell type specificity and distribution of any given lipid species. In conclusion, this large comprehensive dataset of the postmortem human UF phospholipids and cholesterol will serve as a starting point for future research into this unique WM tract and how its molecular properties are impacted in the aging brain.

## Acknowledgements

We would like to thank the technicians of the Douglas-Bell Canada Brain Bank. Finally, we would like to sincerely thank the brain donors and their families for their invaluable gift.

## Data availability statement

Raw chromatograms are available at this link https://osf.io/w97ap/overview?view_only=46295308bde8480589c1e4c7f6ac2ff2 and can be viewed on the CompassCDS software. Western blot and qPCR raw data are available upon request.

## Author contributions

KP: conceptualization, methodology, formal analysis, investigation, writing-original draft; CTC: methodology, investigation, validation, writing-review and editing; MES: methodology, investigation, writing-review and editing; JK: methodology, formal analysis, investigation; GT: resources, writing-review and editing; RPB: conceptualization, project administration, resources, writing-review and editing, supervision; NM: conceptualization, project administration, resources, funding acquisition, writing-review and editing, supervision.

