## Supplementary Material for "A comprehensive characterization of the phospholipid and cholesterol composition of the uncinate fasciculus in the human brain: evidence of age-related alterations"

Supplementary Table 1 – Internal standard information for lipid processing.

| **Internal standard** | **Amount added per sample (ug)** | **Step at which added** |
| --- | --- | --- |
| 5-alpha cholestane (9.95 mg/ml) | 428 | Add to initial homogenate |
| 17:0 PC (1mg/mL) | 61 | Add to initial homogenate |
| 17:0 PE (1mg/mL) | 41 | Add to initial homogenate |
| FFA 17:0 (PtdSer) (0.1mg/mL) | 1.5 | Add to scrapes before methylation |
| FFA 17:0 (PtdIns) (0.1mg/mL) | 1 | Add to scrapes before methylation |
| FFA 17:0 (CerPCho) (0.1mg/mL) | 2 | Add to scrapes before methylation |

Supplementary Table 2 – Primer information for RT-qPCR experiments

| **Gene** | **Forward primer (5’-3’)** | **Reverse primer (5’-3’)** | **Amplicon size (base pairs)** |
| --- | --- | --- | --- |
| ***CNP*** | CATCATGAACAGAGGCTTCTCCC | AACTGCAGCTCAGGCTTGTC | 105 |
| ***GAPDH*** | TTGTCAAGCTCATTTCCTGG | TGTGAGGAGGGGAGATTCAG | 202 |
| ***MAG*** | GCAACCCGGACCCTATTCTC | GGGGCGAACTCCACAGAC | 187 |
| ***MBP*** | TTTAAGCTGGGAGGAAGAGATAGT | GCATTAGGGGAGGGGTCATC | 167 |
| ***MOBP*** | GGCTCTCCAAGAACCAGAAG | GCTTGGAGTTGAGGAAGGTG | 78 |
| ***MOG*** | TGTTGGCCTCATCTTCCTCTG | TGGAGATTCTCTATCTCTGCTCG | 72 |
| ***PLLP*** | CGTTCTCTACATCACCGCCT | ACACGCAAAGAACGAGGCA | 109 |
| ***PLP1*** | TGGTTTCCCTGCTCACCTTC | TGGGAGACGCAGCATTGTAG | 154 |

Supplementary Table 3 – Antibody information for immunoblotting experiments

| **Protein target** | **Manufacturer** | **Species** | **Concentration** |
| --- | --- | --- | --- |
| **MAG** | Abcam | Mouse | 1 in 2500 |
| **MBP** | BioLegend | Mouse | 1 in 10000 |
| **MOBP** | Santa Cruz | Goat | 1 in 4000 |
| **MOG** | Abcam | Rabbit | 1 in 10000 |
| **PLP** | Abcam | Rabbit | 1 in 4000 |

Supplementary Table 4 – Subject information for lipid experiments. Data are demonstrated as mean ± standard error of the mean. P-values are derived from one-way anovas.

|  | **CTRL** | **DS** | **DS-CA** |
| --- | --- | --- | --- |
| **N total** | 20 | 31 | 29 |
| **Male/Female** | 15/5 | 26/5 | 19/10 |
| **Age** (p = 0.025) | 48.70 ± 4.96 | 52.55 ± 1.63 | 41.66 ± 2.76 |
| **pH** (p = 0.69) | 6.45 ± 0.069 | 6.49 ± 0.065 | 6.53 ± 0.057 |
| **PMI (h)** (p = 0.073) | 50.64 ± 5.68 | 67.38 ± 4.73 | 62.78 ± 4.49 |
| **N antidepressants** | 0 | 16 | 13 |

Supplementary Table 5– Subject information for immunoblotting experiments. Data are demonstrated as mean ± standard error of the mean. P-values are derived from one-way anovas.

|  | **CTRL** | **DS** | **DS-CA** |
| --- | --- | --- | --- |
| **N total** | 16 | 23 | 28 |
| **Male/Female** | 15/1 | 20/3 | 25/3 |
| **Age** (p < 0.001) | 38.94 ± 3.87 | 52.57 ± 2.77 | 36.82 ± 2.23 |
| **pH** (p = 0.82) | 6.49 ± 0.066 | 6.46 ± 0.078 | 6.43 ± 0.058 |
| **PMI (h)** (p = 0.028) | 34.32 ± 6.33 | 60.28 ± 6.31 | 51.02 ± 5.68 |
| **N antidepressants** | 1 | 11 | 7 |

Supplementary Table 6– Subject information for RT-qPCR experiments. Data are demonstrated as mean ± standard error of the mean. P-values are derived from one-way anovas.

|  | **CTRL** | **DS** | **DS-CA** |
| --- | --- | --- | --- |
| **N total** | 16 | 24 | 29 |
| **Male/Female** | 15/1 | 21/3 | 26/3 |
| **Age** (p < 0.001) | 38.13 ± 3.73 | 53.58 ± 2.85 | 36.59 ± 2.16 |
| **pH** (p = 0.85) | 6.50 ± 0.067 | 6.46 ± 0.075 | 6.44 ± 0.057 |
| **PMI (h)** (p = 0.072) | 36.95 ± 6.51 | 58.88 ± 6.20 | 51.04 ± 5.48 |
| **N antidepressants** | 1 | 11 | 7 |

Supplementary Table 7– FA special metrics and indices for all phospholipid fractions and the TL. Data are demonstrated as mean ± standard deviation. *Not calculated due to low concentration of omega-3 FAs.

|  | **ChoGpl** | **EtnGpl** | **CerPCho** | **PtdSer** | **PtdIns** | **TL** |
| --- | --- | --- | --- | --- | --- | --- |
| **Mean Chain Length Index** | 1740.71 ± 2.97 | 1959.14 ± 11.97 | 2072.01 ± 52.57 | 1854.11 ± 18.94 | 1834.25 ± 26.36 | 1876.13 ± 36.09 |
| **Mean Unsaturation Index** | 70.82 ± 2.42 | 242.44 ± 17.40 | 40.17 ± 6.59 | 106.59 ± 24.09 | 128.87 ± 22.11 | 135.01 ± 15.98 |
| **Mean Peroxidation Index** | 29.69 ± 6.06 | 227.17 ± 33.54 | 2.51 ± 0.54 | 78.11 ± 36.97 | 105.54 ± 25.13 | 108.43 ± 19.20 |
| **Mean Omega-6/Omega-3 Ratio** | 5.52 ± 0.82 | 3.89 ± 1.35 | -* | 2.00 ± 2.53 | 11.12 ± 4.52 | 4.10 ± 2.21 |
| **Mean HUFA (µg/g)** | 577.78 ± 109.44 | 4612.50 ± 660.04 | 6.56 ± 2.83 | 730.19 ± 217.21 | 153.93 ± 108.78 | 7498.34 ± 2114.21 |

Supplementary Table 8 – FA concentrations of ChoGpl for the UF and the ACC. Data are demonstrated as mean ± standard deviation.

| **Fatty acid** | **UF ChoGpl (µg/g)** | **ACC ChoGpl (µg/g)** |
| --- | --- | --- |
| **C 14:0** | 85.49 ± 13.47 | 86.96 ± 17.51 |
| **C 16:0; PAM** | 3490.12 ± 424.73 | 3397.46 ± 614.38 |
| **C 18:0; STA** | 1206.27 ± 158.77 | 1401.18 ± 248.57 |
| **C 20:0** | 7.89 ± 1.58 | 10.44 ± 2.40 |
| **C 22:0** | 2.43 ± 0.77 | 9.23 ± 2.77 |
| **C 24:0** | 7.13 ± 4.48 | 16.11 ± 6.85 |
| **C 16:1n-7** | 121.88 ± 23.82 | 124.73 ± 22.35 |
| **C 18:1n-7** | 628.78 ± 95.08 | 747.65 ± 122.35 |
| **C 18:1n-9 (OLA)** | 3400.93 ± 578.75 | 4066.80 ± 690.47 |
| **C 20:1n-9** | 69.11 ± 16.80 | 97.99 ± 24.52 |
| **C 22:1n-9** | 10.62 ± 4.30 | 10.24 ± 3.76 |
| **C 24:1n-9** | 11.19 ± 3.52 | 24.69 ± 8.95 |
| **C 18:2n-6; LNA** | 64.89 ± 15.62 | 66.12 ± 17.59 |
| **C 20:2n-6** | 8.79 ± 10.10 | 10.84 ± 3.13 |
| **C 20:3n-6; DGLA** | 48.33 ± 11.36 | 43.58 ± 11.78 |
| **C 20:4n-6; ARA** | 333.40 ± 65.70 | 233.76 ± 51.65 |
| **C 22:4n-6; AdA** | 74.19 ± 11.77 | 67.43 ± 16.14 |
| **C 22:5n-6; DPAn-6** | 10.31 ± 3.77 | 9.68 ± 4.77 |
| **C 22:6n-3; DHA** | 95.30 ± 25.07 | 66.25 ± 19.41 |
| **Total FA** | 9689.42 ± 1219.34 | 10566.85 ± 1754.43 |

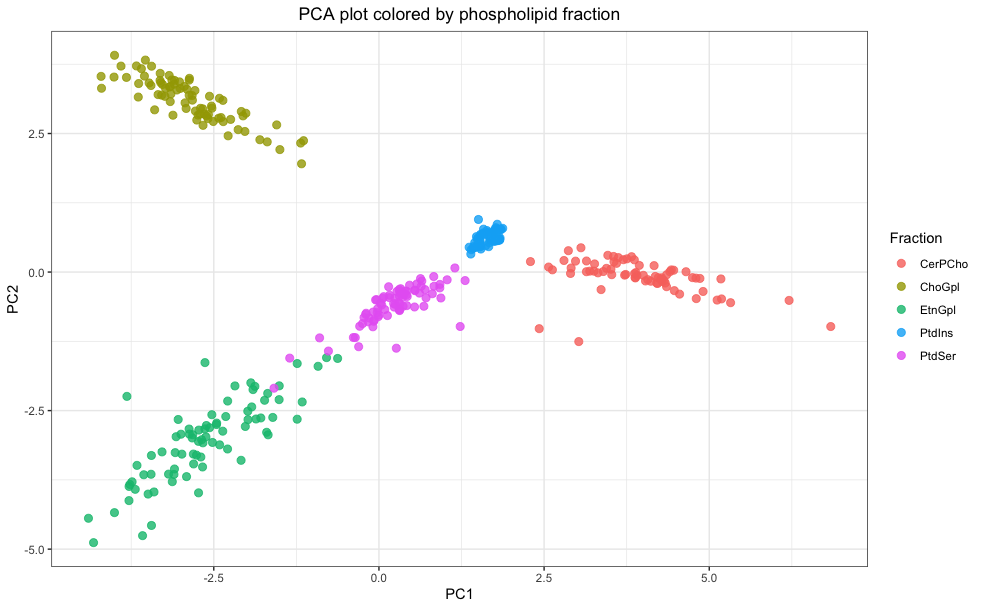

Supplementary Figure 1 – PCA plot colored by phospholipid fractions. The x-axis represents the first principal component, and the y-axis represents the second principal component.

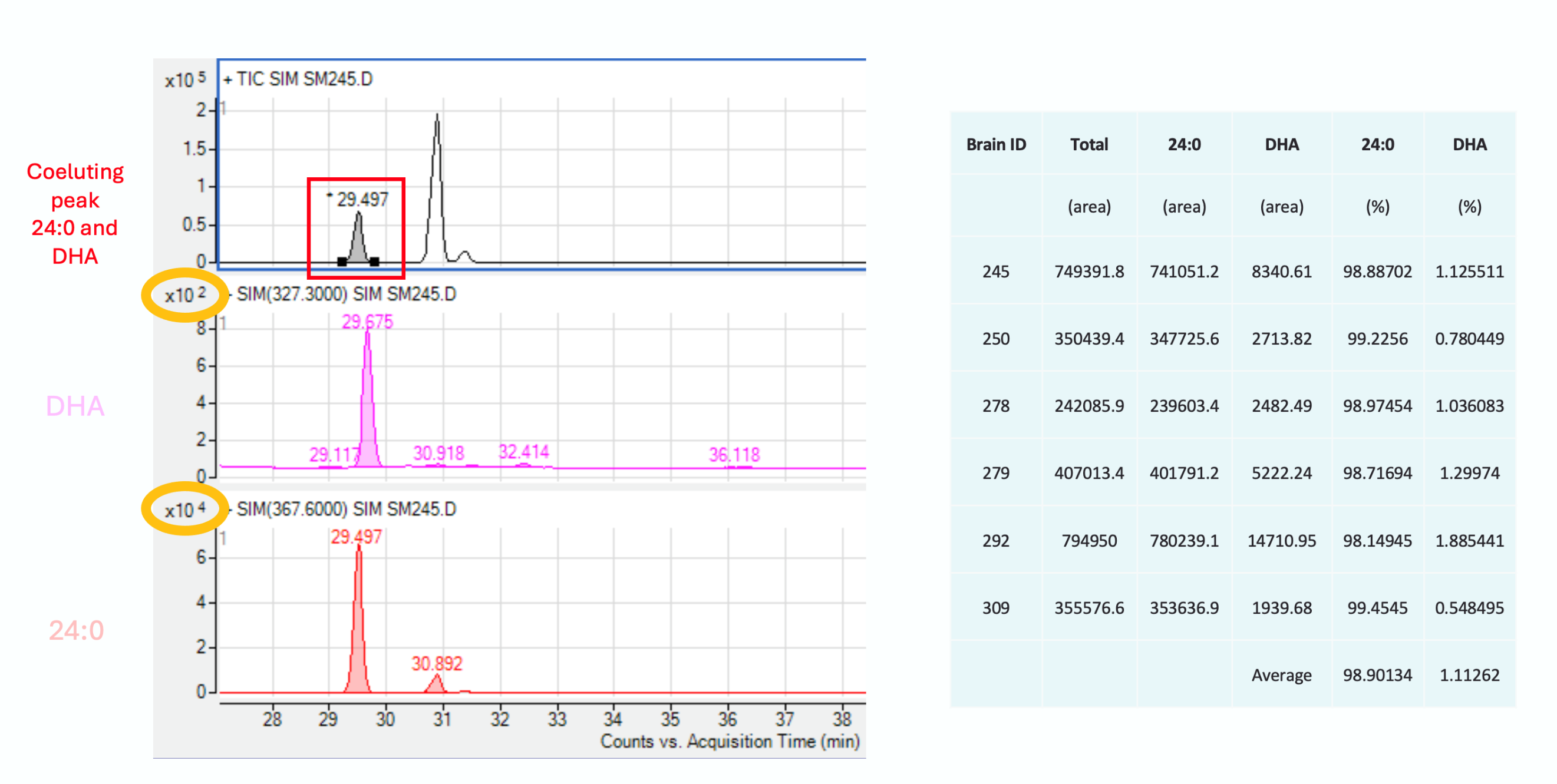

**B**

**A**

Supplementary Figure 2 – Sphingomyelin (CerPCho) co-elution of DHA and C24:0 validation performed by gas chromatography-mass spectrometry (GC-MS) analysis. Supplementary analysis to determine the relative contribution of the co-eluting peak between C24:0 and DHA (C22:6n-3). A) Chromatograms for the co-eluting peak (top), DHA (middle), and C24:0 (bottom). The yellow circles highlight the scale of the y-axis, demonstrating the comparative abundance of C24:0 compared to DHA in CerPCho. B) Table with 6 test subjects with area and percentage values for C24:0 and DHA. On average, the co-eluting peak is composed of 98.9% C24:0 and 1.1% DHA.

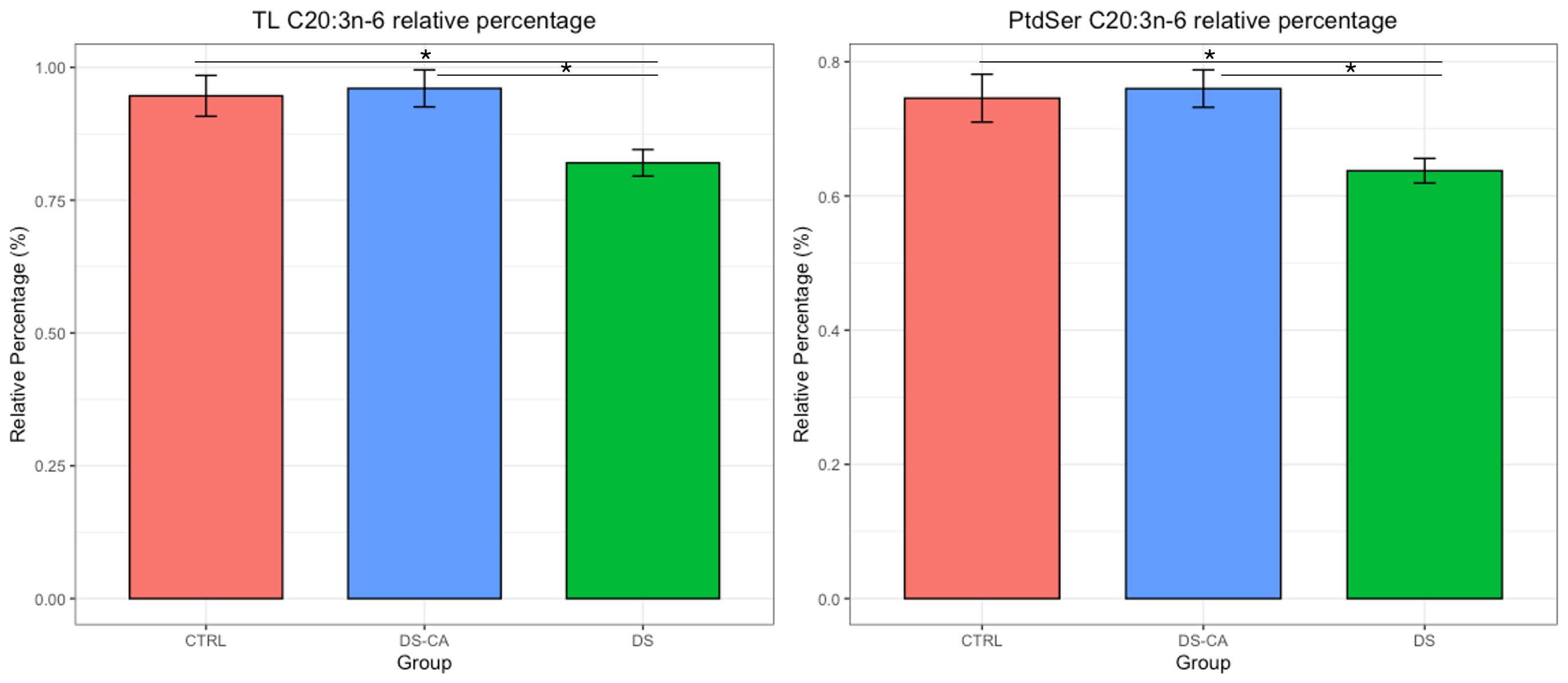

Supplementary Figure 3 – Bar plots for FA quantities that show nominally significance between groups. Significance stars indicate nominal p < 0.05. Data are represented as mean ± standard error. None of these FA retain significance after correction for multiple comparisons.

**A**

**B**

ChoGpl relative percent

ChoGpl concentration

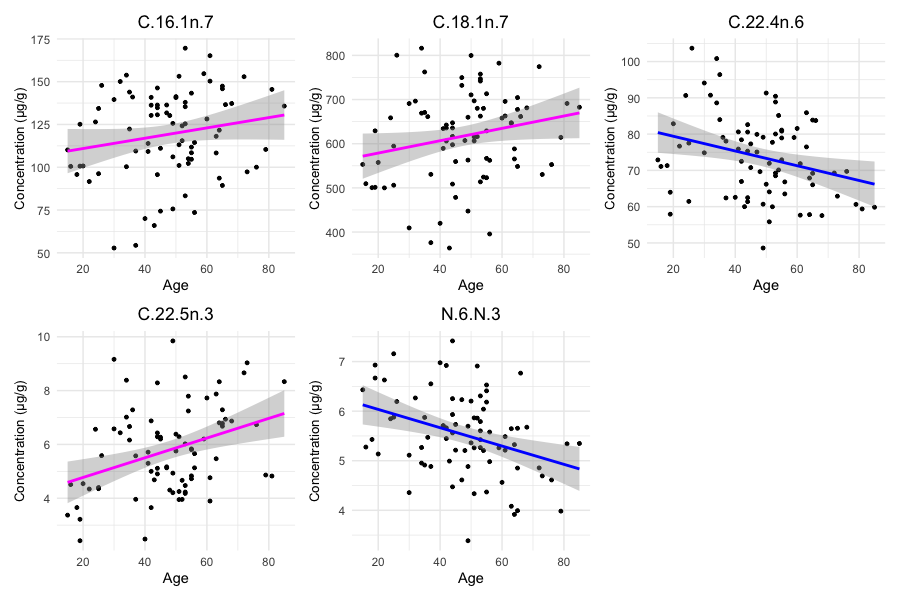

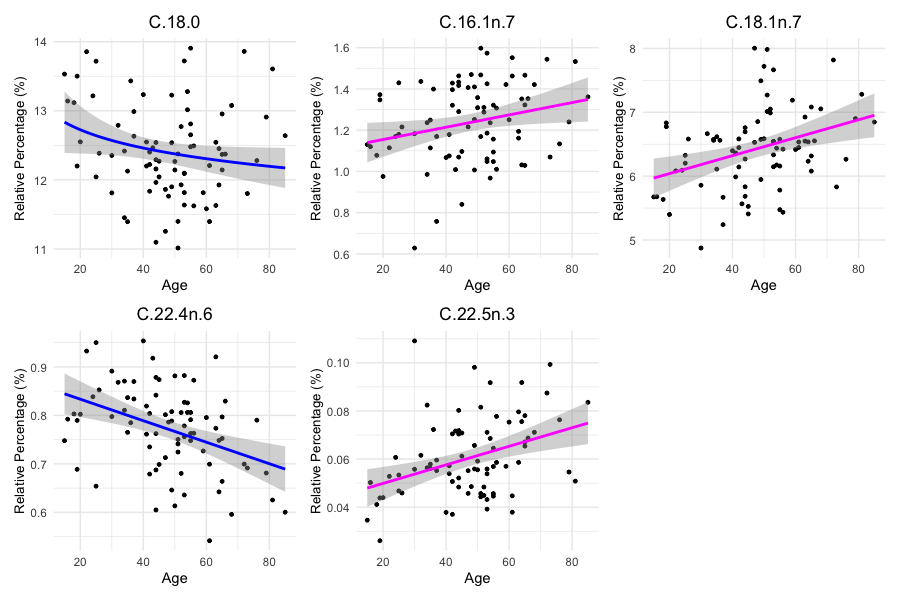

Supplementary Figure 4 – Scatter plots with regression line showing significant relationships between age and ChoGpl A) concentration and B) relative percentage. A blue regression line indicates the model coefficient for age is negative, and a magenta line indicates the model coefficient for age is positive.

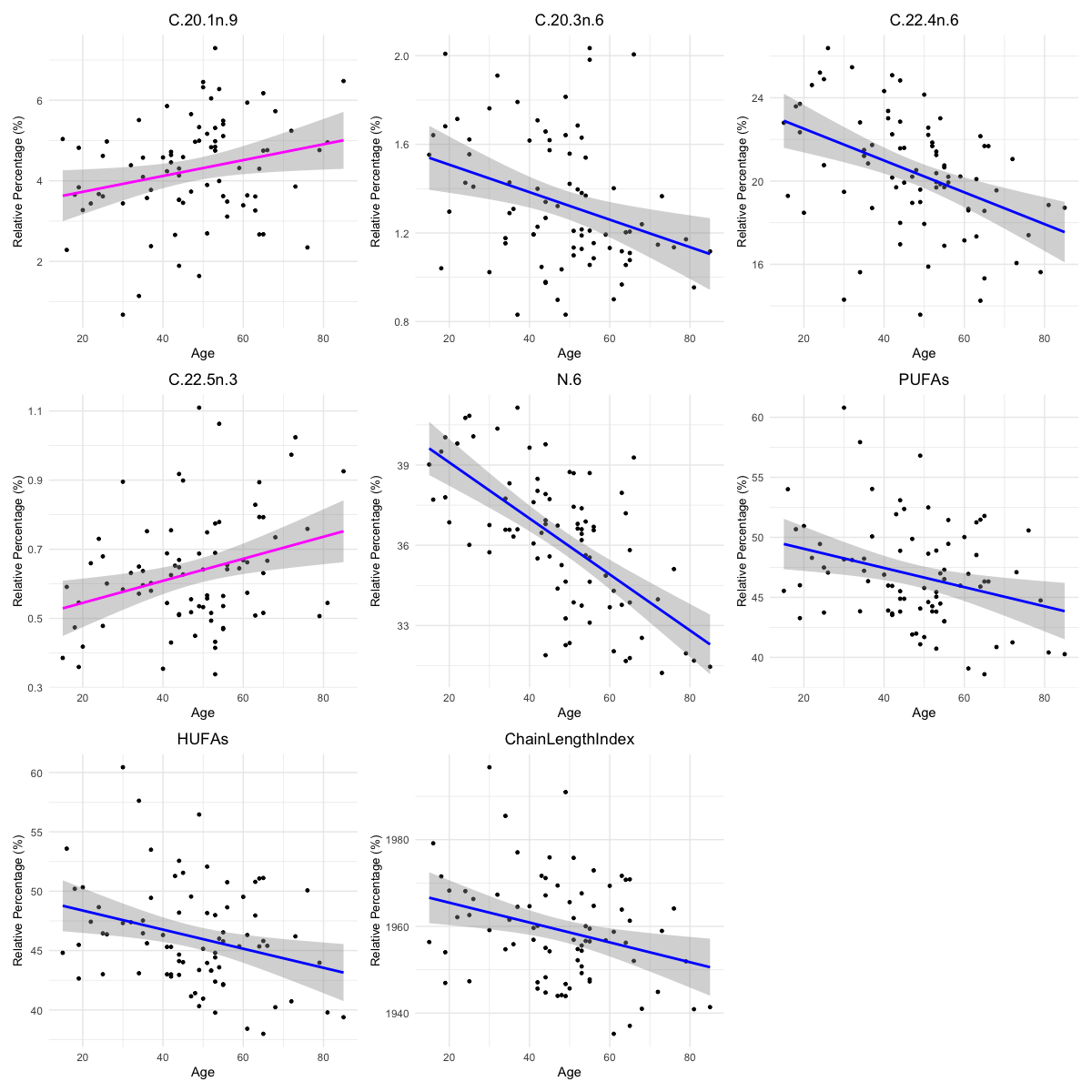

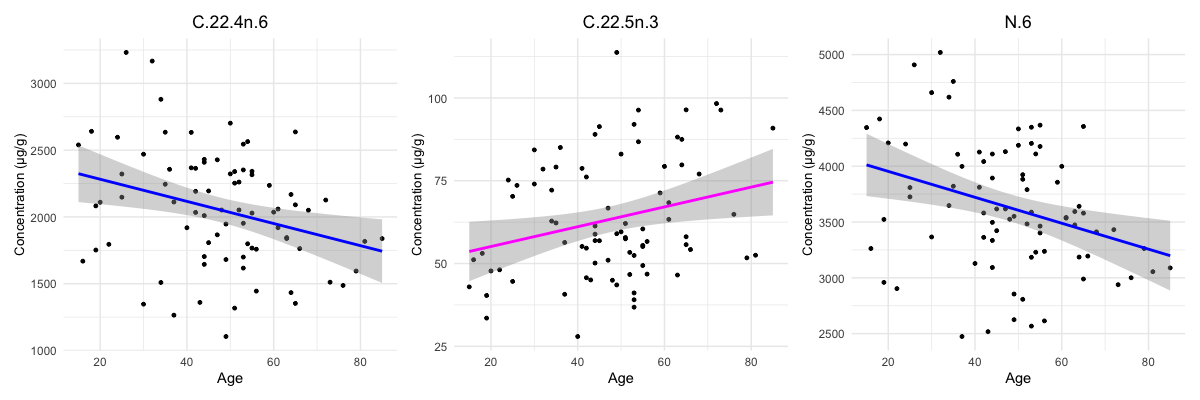

**A**

**B**

EtnGpl relative percent

EtnGpl concentration

Supplementary Figure 5 – Scatter plots with regression line showing significant relationships between age and EtnGpl A) concentration and B) relative percentage. A blue regression line indicates the model coefficient for age is negative, and a magenta line indicates the model coefficient for age is positive.

**A**

**B**

PtdSer concentration

PtdSer relative percent

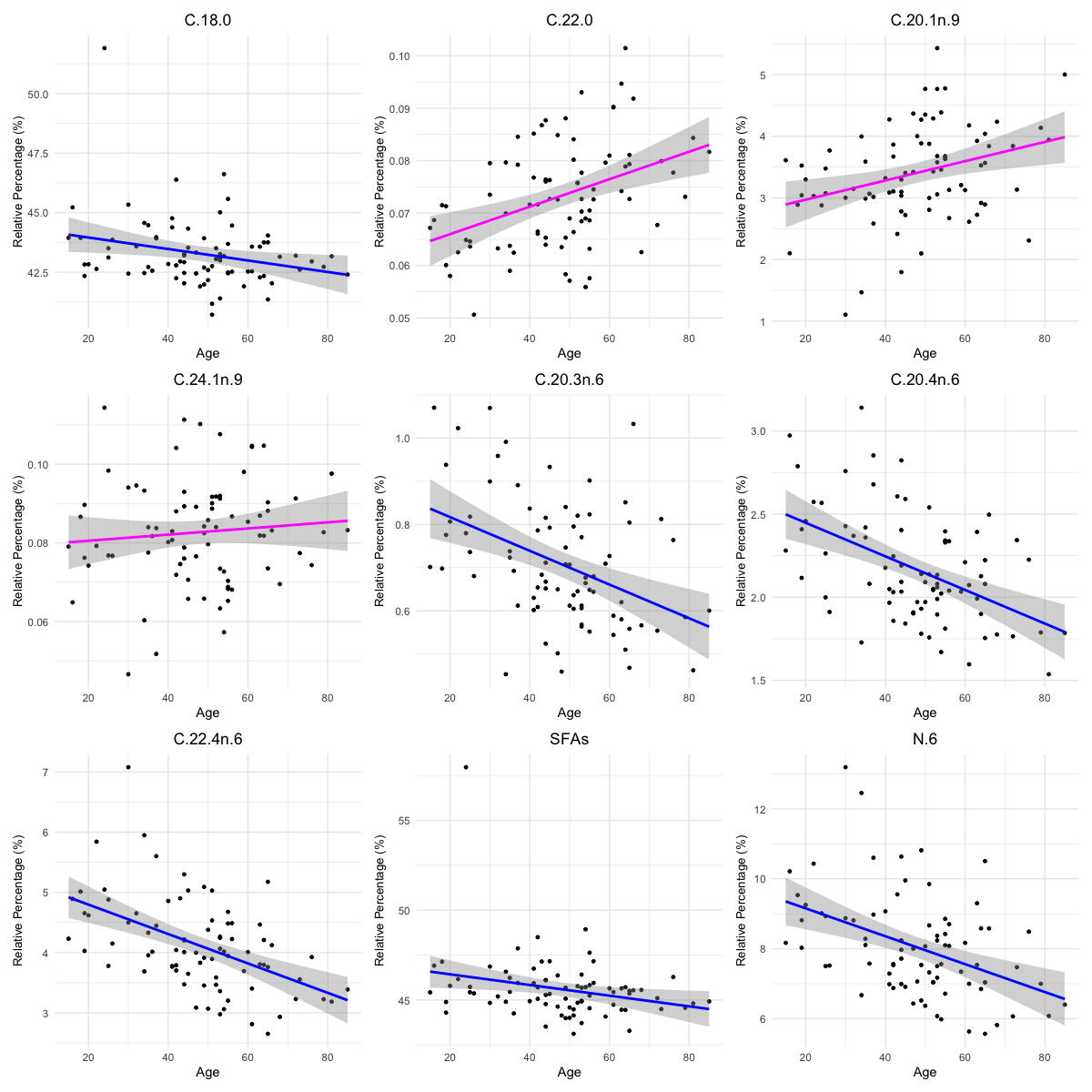

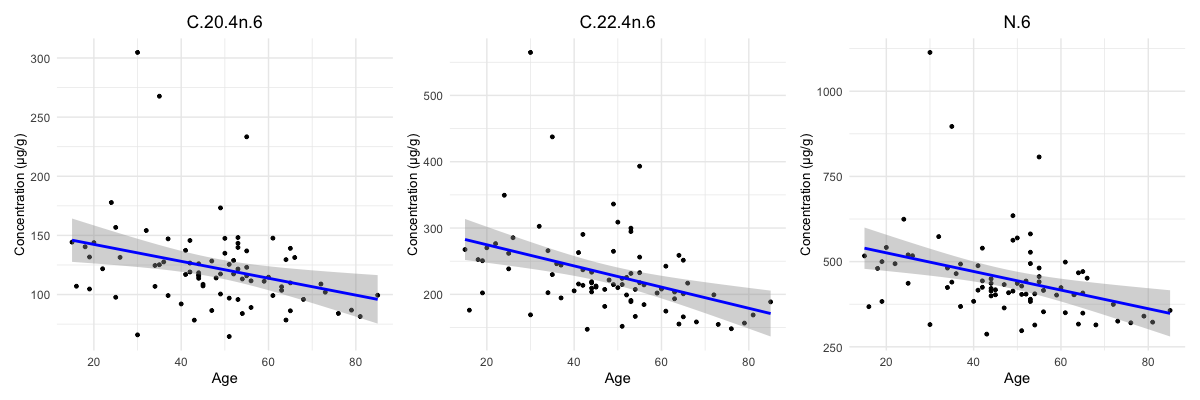

Supplementary Figure 6– Scatter plots with regression line showing significant relationships between age and PtdSer A) concentration and B) relative percentage. A blue regression line indicates the model coefficient for age is negative, and a magenta line indicates the model coefficient for age is positive.

**B**

**A**

PtdIns relative percent

PtdIns concentration

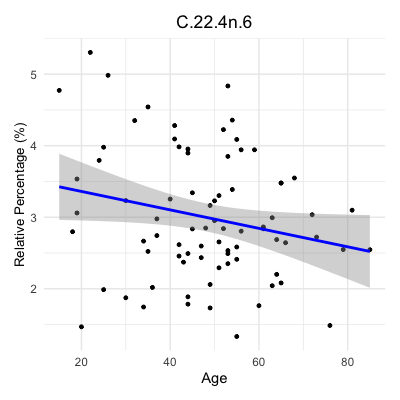

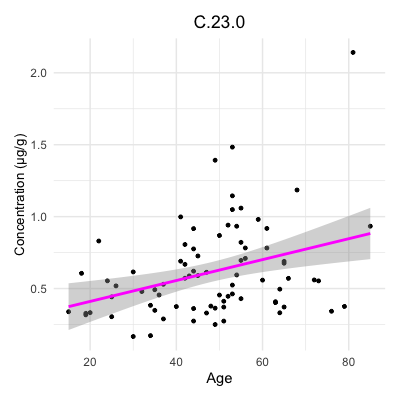

Supplementary Figure 7 – Scatter plots with regression line showing significant relationships between age and PtdIns A) concentration and B) relative percentage. A blue regression line indicates the model coefficient for age is negative, and a magenta line indicates the model coefficient for age is positive.

**B**

**A**

TL concentration

TL relative percent

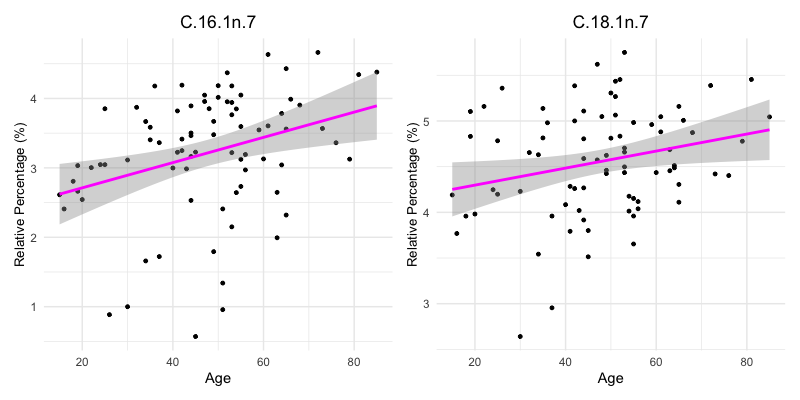

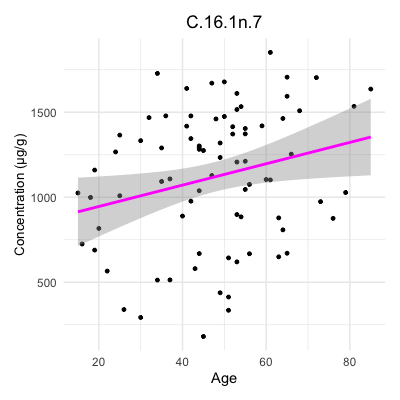

Supplementary Figure 8– Scatter plots with regression line showing significant relationships between age and TL A) concentration and B) relative percentage. A magenta line indicates the model coefficient for age is positive.

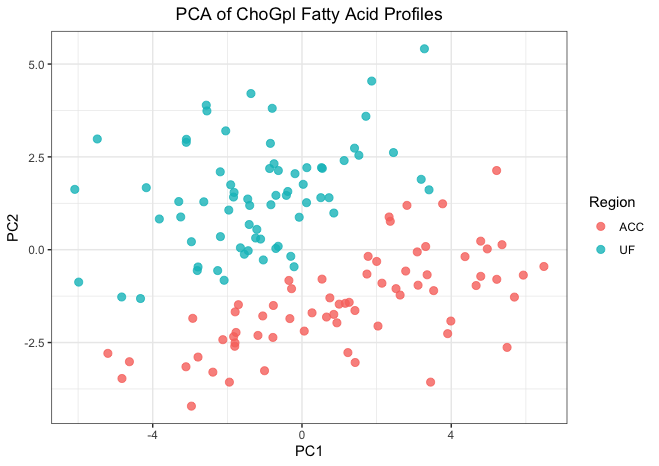

Supplementary Figure 9 - PCA plot of ChoGpl FA profiles colored by region. The x-axis represents the first principal component, and the y-axis represents the second principal component. Dots in pink correspond to the ACC and dots in teal correspond to UF.

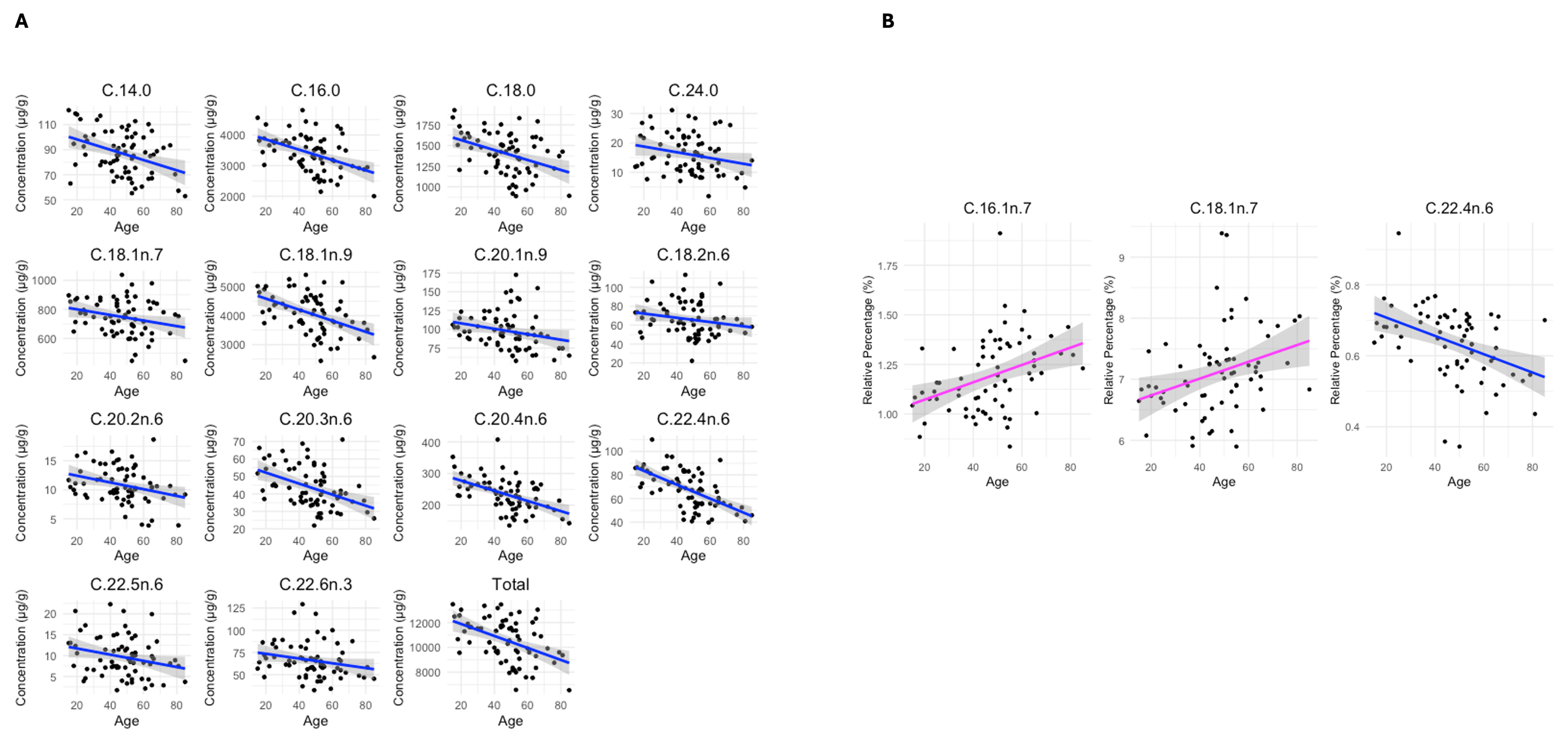

Supplementary Figure 10 – Scatter plots with regression line showing significant relationships between age and ACC A) concentration and B) relative percentage. A blue regression line indicates the model coefficient for age is negative, and a magenta line indicates the model coefficient for age is positive. All significant models were best fit by a linear age term. Nominal p-values for the plotted FAs: C14:0 concentration p = 0.0008; C16:0 concentration p = 0.0011; C 18:0 concentration p=0.0001; C 24:0 concentration p = 0.033; C 16:1n-7 relative percentage p = 0.0002; C18:1n-7 concentration p=0.045, relative percentage p=0.022; C18:1n-9 concentration p = 0.0008; C 20:1n-9 concentration p=0.045, C18:2n-6 concentration p = 0.020; C20:2n-6 concentration p = 0.018; C20:3n-6 concentration p = 0.0011; C20:4n-6 concentration p = 0.0001; C22:4n-6 concentration p < 0.00001, relative percentage p = 0.0008; C22:5n-6 concentration p =0.032; C22:6n-3 concentration p =0.047; total concentration p =0.0005. All of these values survive correction for multiple comparisons (BH-corrected value p < 0.05) except for C18:1n-7 relative percentage, C22:5n-6 concentration, C24:0 concentration, C 18:1n-7 concentration, C20:1n-9 concentration, and C22:6n-3 concentration.

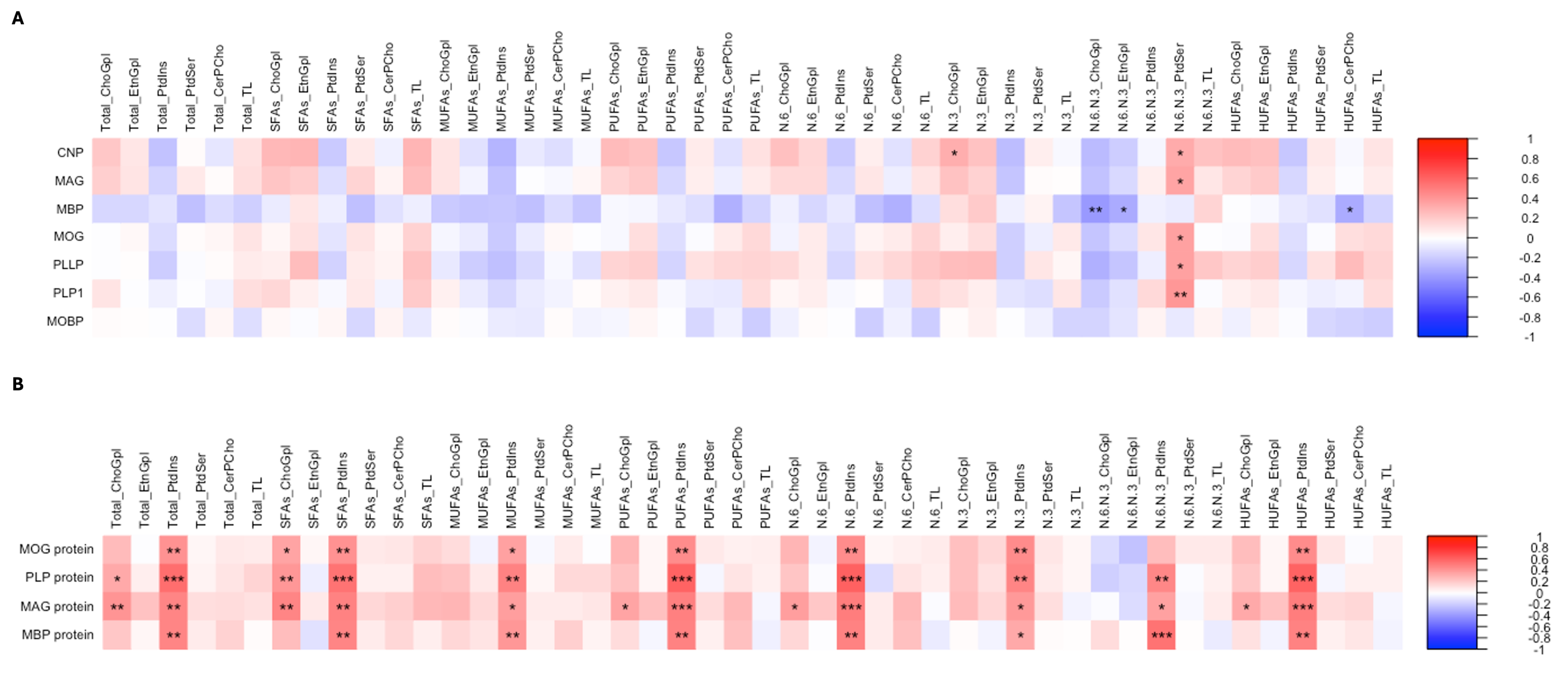

Supplementary Figure 11 - Correlation plots showing the relationships between summary concentration metrics (ug/g) for all FA classes and myelin-constituent genes (A) and proteins (B). Due to the exploratory nature of this analysis, p-values were not corrected for multiple comparisons. The color bar represents Pearson’s correlation coefficient between -1 and 1, where negative coefficients are blue in color and positive coefficients are red in color. *** = p < 0.001, ** = p < 0.01,* = p <0.05

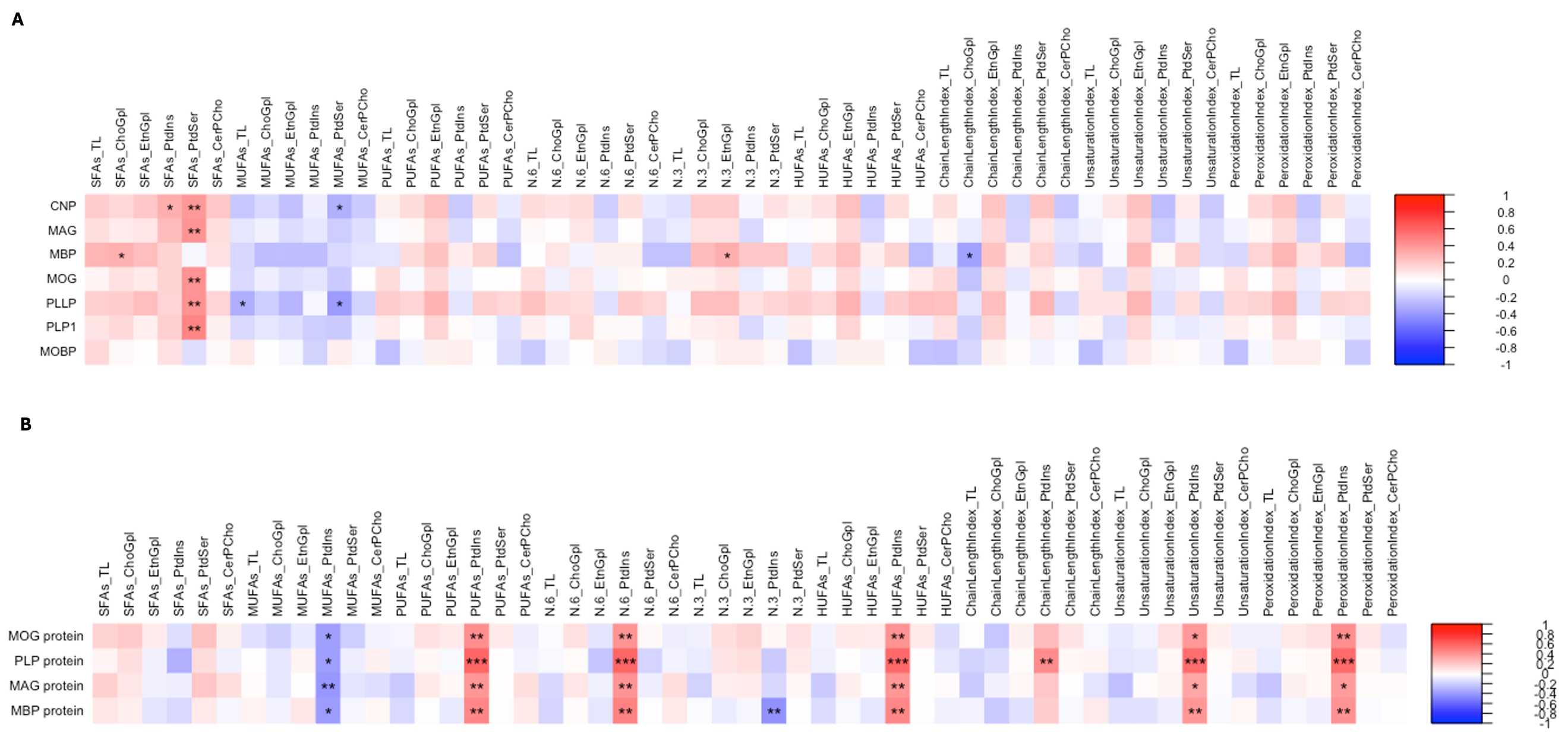

Supplementary Figure 12 - Correlation plots showing the relationships between summary relative percentage metrics (%) for all FA classes and myelin-constituent genes (A) and proteins (B). Due to the exploratory nature of this analysis, p-values were not corrected for multiple comparisons. The color bar represents Pearson’s correlation coefficient between -1 and 1, where negative coefficients are blue in color and positive coefficients are red in color. *** = p < 0.001, ** = p < 0.01,* = p <0.05
